# CIFR (*Clone–Integrate–Flip-out–Repeat*): a toolset for iterative genome and pathway engineering of Gram-negative bacteria

**DOI:** 10.1101/2024.11.07.622458

**Authors:** Filippo Federici, Francesco Luppino, Clara Aguilar-Vilar, Maria Eleni Mazaraki, Lars Boje Petersen, Linda Ahonen, Pablo I. Nikel

## Abstract

Advances in genome engineering enable precise and customizable modifications of bacterial species. Toolsets for metabolic engineering that exhibit broad-host compatibility are particularly valued owing to their portability. Tn*5* transposon vectors have been widely used to establish random integrations of desired DNA sequences into bacterial genomes. However, the iteration of the procedure remains challenging because of the limited availability of selection markers. Here, we present CIFR, a mini-Tn*5* integration system for iterative genome engineering. The pCIFR vectors incorporate *attP* and *attB* sites flanking an antibiotic resistance marker used to select for the insertion. Subsequent removal of these antibiotic determinants is facilitated by the Bxb1 integrase, and a user-friendly counter-selection marker, both encoded in auxiliary plasmids. Hence, CIFR delivers engineered strains harboring stable DNA insertions and free of any antibiotic resistance cassette, allowing for the reusability of the tool. The system was validated in *Pseudomonas putida, Escherichia coli*, and *Cupriavidus necator*, underscoring its portability across diverse industrially-relevant hosts. The CIFR toolbox was calibrated through combinatorial integrations of chromoprotein genes in *P. putida*, generating strains displaying a diverse color *palette*. Next, we introduced a carotenoid biosynthesis pathway in *P. putida* in a two-step engineering process, showcasing the potential of the tool for pathway balancing. The broad utility of the toolbox expands the synthetic biology toolkit for metabolic engineering, allowing for the construction of complex phenotypes, while opening new possibilities in bacterial genetic manipulations.

## 1. INTRODUCTION

The challenges of modern society—ranging from climate change and pollution to the need for sustainable material production—are driving bio-based solutions to the forefront of metabolic engineering research. Engineered microbes are increasingly being used in diverse applications, from CO_2_ assimilation and upcycling to the production of bulk chemicals and pharmaceuticals [1, 2]. Over the years, advances in genome engineering tools have greatly expanded our ability to reprogram both model and non-canonical bacterial species for industrial purposes [3–5]. However, the full potential of microbial biotechnology can only be realized with toolsets that enable stable, efficient, and controllable gene expression across a variety of bacterial species and applications.

Achieving stable and balanced expression of multi-gene pathways remains a substantial challenge in metabolic engineering [6, 7]. Microbial biotechnology relies on the heterologous expression of multiple genes that must be finely tuned for optimal performance. Appropriate pathway balancing is critical to avoid metabolic bottlenecks, toxic intermediate accumulation, or resource misallocation, which ultimately reduce titer, rates, and yield [TRY] [8–10]. Traditional approaches to pathway balancing often involve using multiple plasmids, but this strategy leads to phenotypic instability due to plasmid loss, variable copy numbers, and increased metabolic burden on the host [11, 12]. These challenges are particularly relevant in industrial settings, where long-term cultivation is commonplace [13]. Hence, stable genomic integration of target pathways offers a solution by ensuring consistent gene expression across generations while circumventing the reliance on plasmids [14].

Bacterial transposons, discovered around 50 years ago [15], have become valuable tools for metabolic engineering. These mobile genetic elements can integrate large DNA fragments into target genomes, facilitating stable insertion of heterologous pathways. Among these, Tn*5* is a well-known composite transposon encompassing a central region flanked by inverted repeats (IS*50* elements; [16]. The first tools for bacterial genome engineering based on Tn*5*, published by de Lorenzo, Herrero [17] and Herrero, de Lorenzo [18], have been widely adopted for both fundamental and applied research in microbial sciences. Over the years, numerous modifications have been introduced to Tn*5*-based tools, adapting them to different formats and applications (**Table 1**). However, most transposon-based methods still face limitations, e.g., the challenge of reusing functional elements together with antibiotic resistance (Ab^R^) markers that are left in the genome after integration [19, 20]. These limitations pose risks in applications in food or pharma, where the presence of Ab^R^ genes can raise safety concerns. Additionally, retaining resistance markers hinders the iterative engineering of strains [21], as further modifications add to the genetic load. While some tools allow for the excision of Ab^R^ markers post-integration, they often require complex and time-consuming procedures [22], reducing the efficiency and throughput of strain development. One promising solution is the site-specific BxbI phage integrase (BxbI-int) that mediates recombination between two specific sequences (i.e., *attachment phage*, or *attP*, and *attachment bacteria*, or *attB*) and enables precise DNA integration or excision [23]. BxbI-int has been used in microbial, plant, and animal cells for targeted genome modifications due to its efficiency and minimal off-target effects [24–28].

**Table 1.**
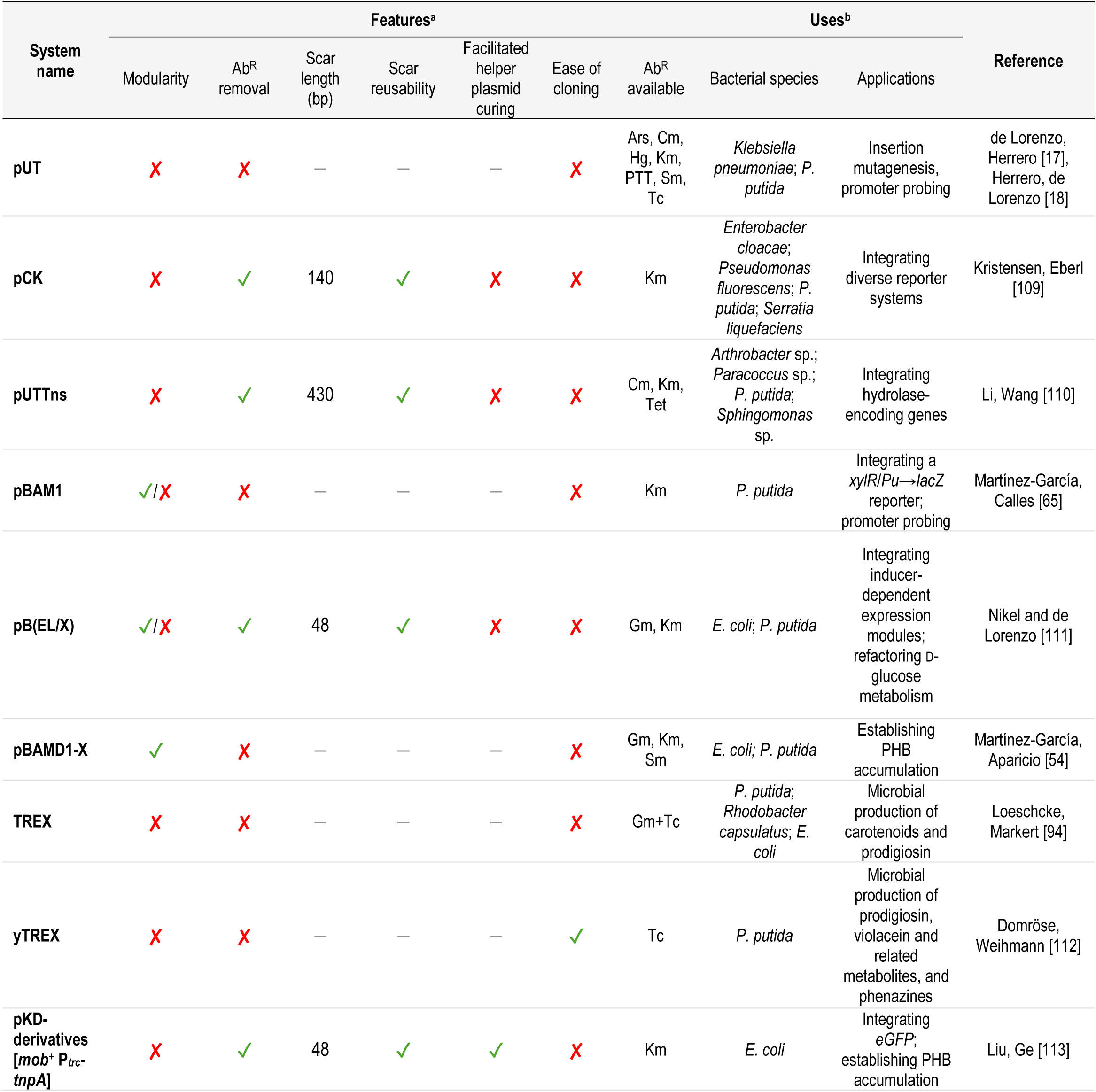
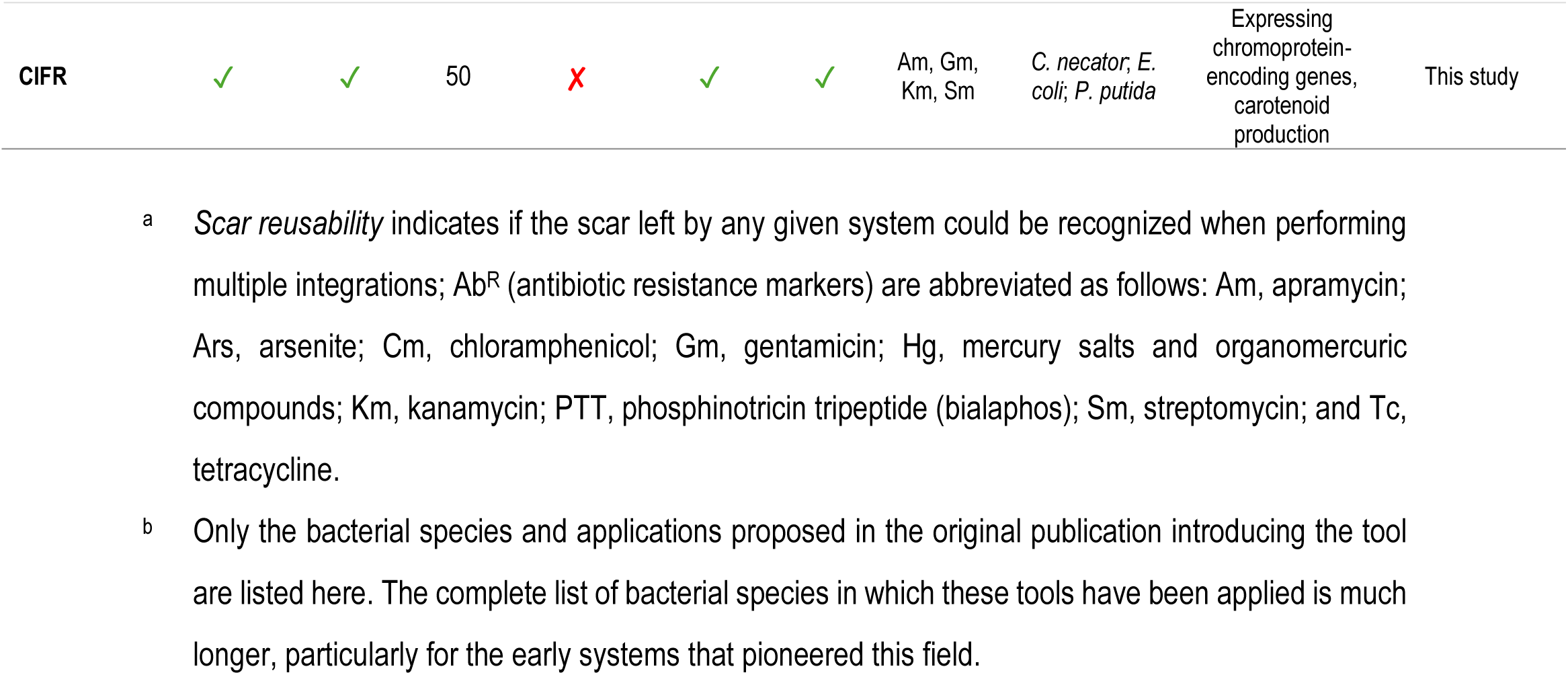
Overview of Tn*5*-based tools applied for genomic integration of synthetic modules in Gram-negative bacteria.

Building on this background, here we introduce the CIFR (*Clone–Integrate–Flip-out–Repeat*) system for efficient genome engineering for Gram-negative bacteria. The CIFR toolset combines features of Tn*5*-mediated integration with the BxbI phage integrase. First, a synthetic mini-Tn*5* transposon is used to stably integrate a defined DNA sequence, including an Ab^R^ marker, into the bacterial genome. Once the desired integration is achieved, BxbI-int is applied to excise the Ab^R^ module, leaving behind only the desired DNA cargo. This process enables multiple rounds of strain engineering without the accumulation of unwanted genetic material, streamlining the development of complex, modularized metabolic pathways. We selected *Pseudomonas putida*, *Escherichia coli*, and *Cupriavidus necator* as the microbial hosts to test the CIFR system. *P*. *putida* is a versatile soil bacterium, capable of degrading environmental pollutants and synthesizing a wide range of valuable chemicals, while exhibiting tolerance under industrial conditions [29–31]. *E*. *coli* remains one of the most widely studied model bacteria, frequently used in the production of recombinant proteins and multiple bio-based chemicals [32]. *C*. *necator* is widely known for its ability to convert CO_2_ into polyhydroxyalkanoates and other products, making it an attractive platform for producing sustainable chemicals from renewable carbon sources [33].

To validate the CIFR system, we applied this toolset to engineer stable production of chromoproteins (CPs) and to establish carotenoid biosynthesis, which serve as ideal testbeds for pathway engineering.

CPs are often employed as reporter genes, and having a variety of alternative CPs with different colors and properties is advantageous for multi-channel imaging and other applications [34]. Carotenoids are naturally occurring pigments with applications in the food, cosmetic, and pharmaceutical industries [35]. Their complex biosynthesis pathways are well-suited for modularization, allowing for precise control over each enzymatic step. By modularizing these reporter modules and biosynthetic pathways using our toolset, we showcase the flexibility and versatility of CIFR in a variety of microbial hosts. The ability to integrate and excise genes with precision and efficiency facilitates stable, iterative pathway development in bacteria. This highlights the potential for broader applications in microbial biotechnology, from industrial bioproduction to environmental remediation.

## 2. MATERIALS AND METHODS

### 2.1. Bacterial strains and culture conditions

The bacterial strains used in this study are listed in **Table 2**. *E. coli* was incubated at 37 °C, *P. putida* and *C. necator* were incubated at 30°C. For propagation and storage, routine cloning procedures, and during genome engineering manipulations [36–40], cells were grown in lysogeny broth (LB) medium (10 g L^−1^ tryptone, 5 g L^−1^ yeast extract, and 10 g L^−1^ NaCl). Liquid cultures were performed in 15-mL conical centrifuge tubes with a medium volume of 2-6 mL. All liquid cultures were agitated at 250 rpm in a MaxQ™8000 incubator (Thermo Fisher Scientific Co., Waltham, MA, USA). Solid culture media contained 15 g L^−1^ agar. For *E. coli* and *P. putida* strains, selection of plasmid-harboring cells or cells with integrated Ab^R^ was achieved by adding kanamycin (Km), streptomycin (Sm), gentamicin (Gm), apramycin (Am), or sucrose to the culture media at 50 μg mL^−1^, 100 μg mL^−1^, 10 μg mL^−1^, 25 μg mL^−1^, and 100 mg mL^−1^, respectively. For *C. necator* strains, selection was achieved by adding chloramphenicol (Cm), Km, Sm, tetracycline (Tc), or sucrose to the culture media at 50 μg mL^−1^, 100 μg mL^−1^, 200 μg mL^−1^, 15 μg mL^−1^, and 100 mg mL^−1^, respectively.

**Table 2.**
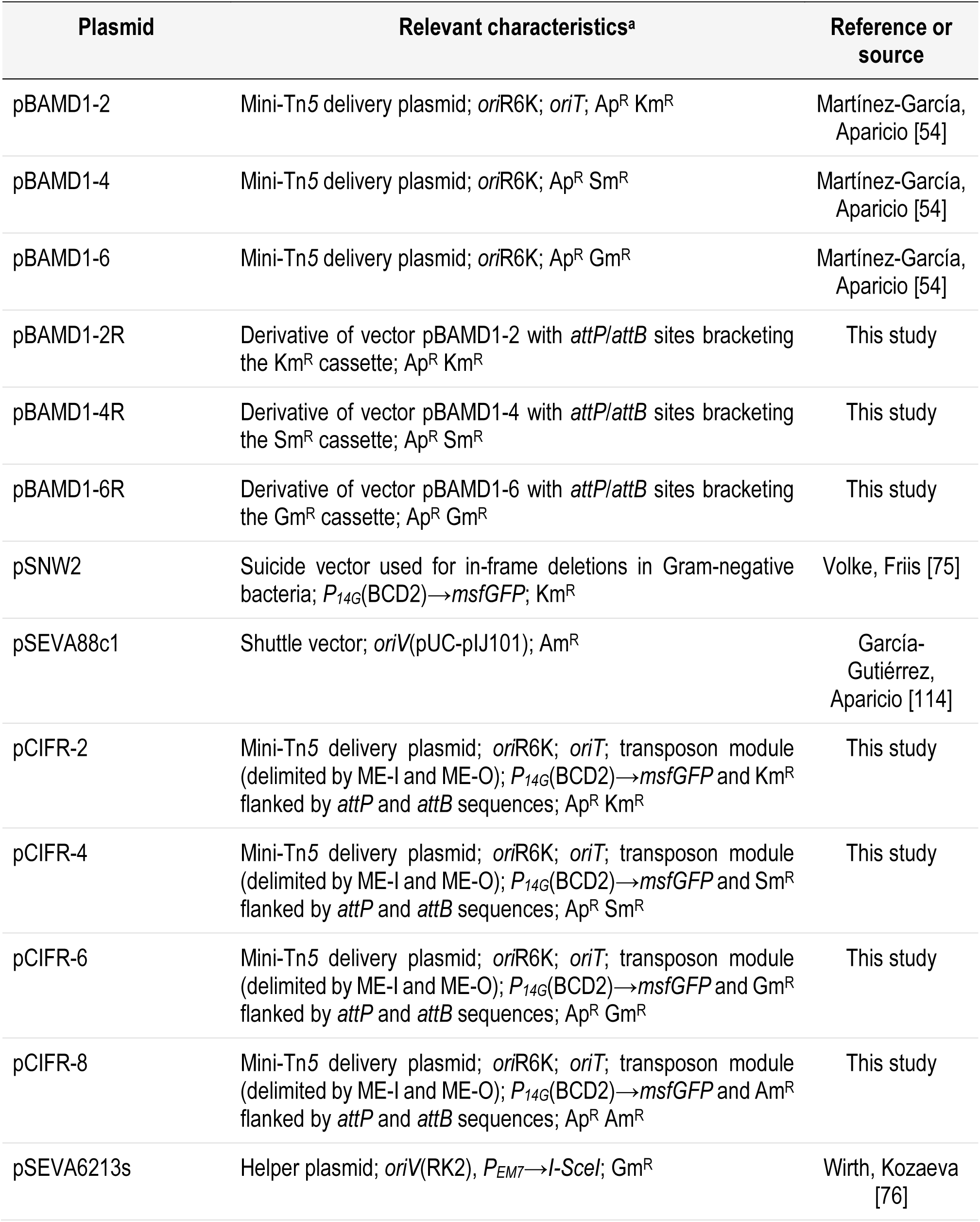
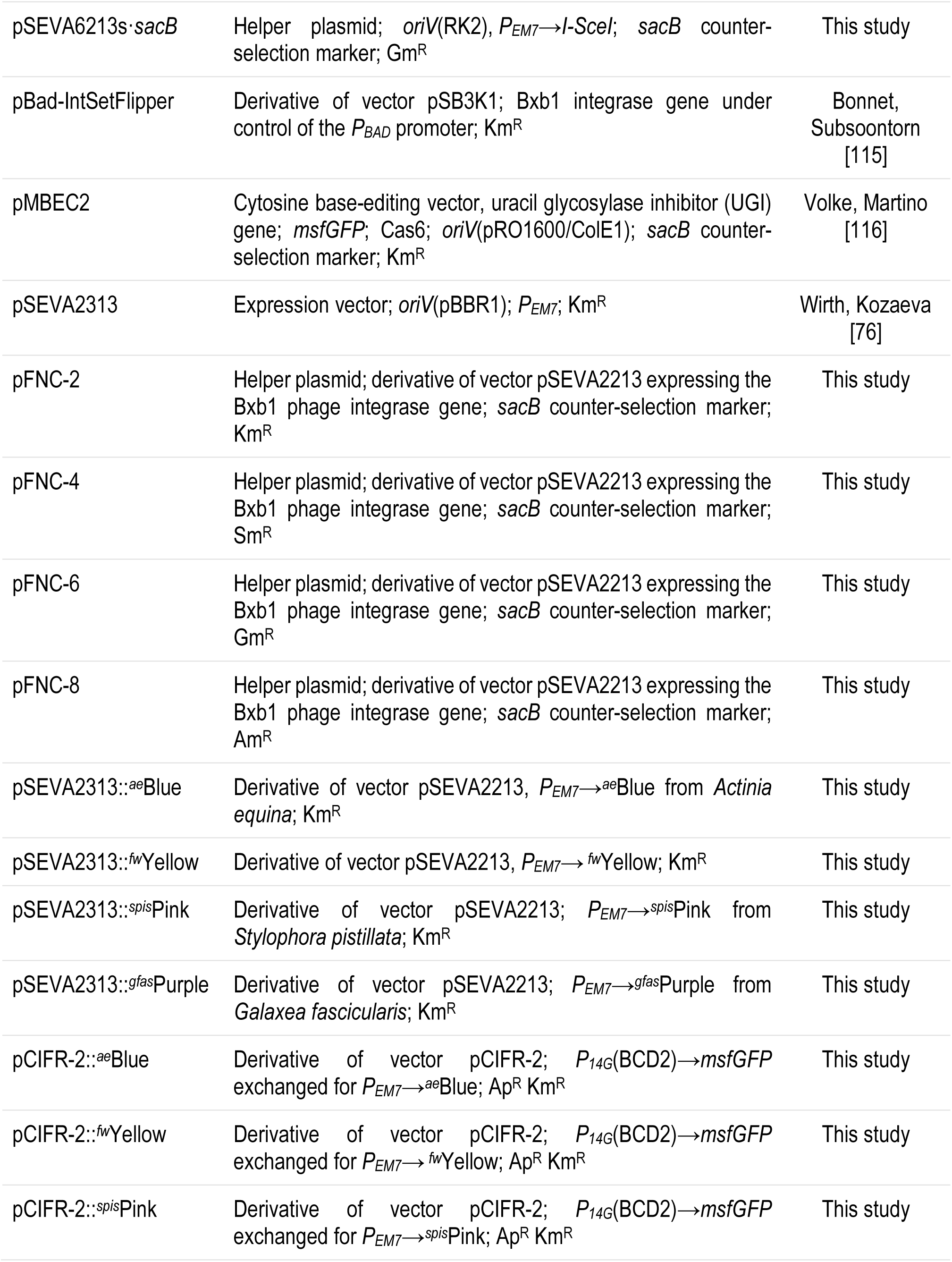
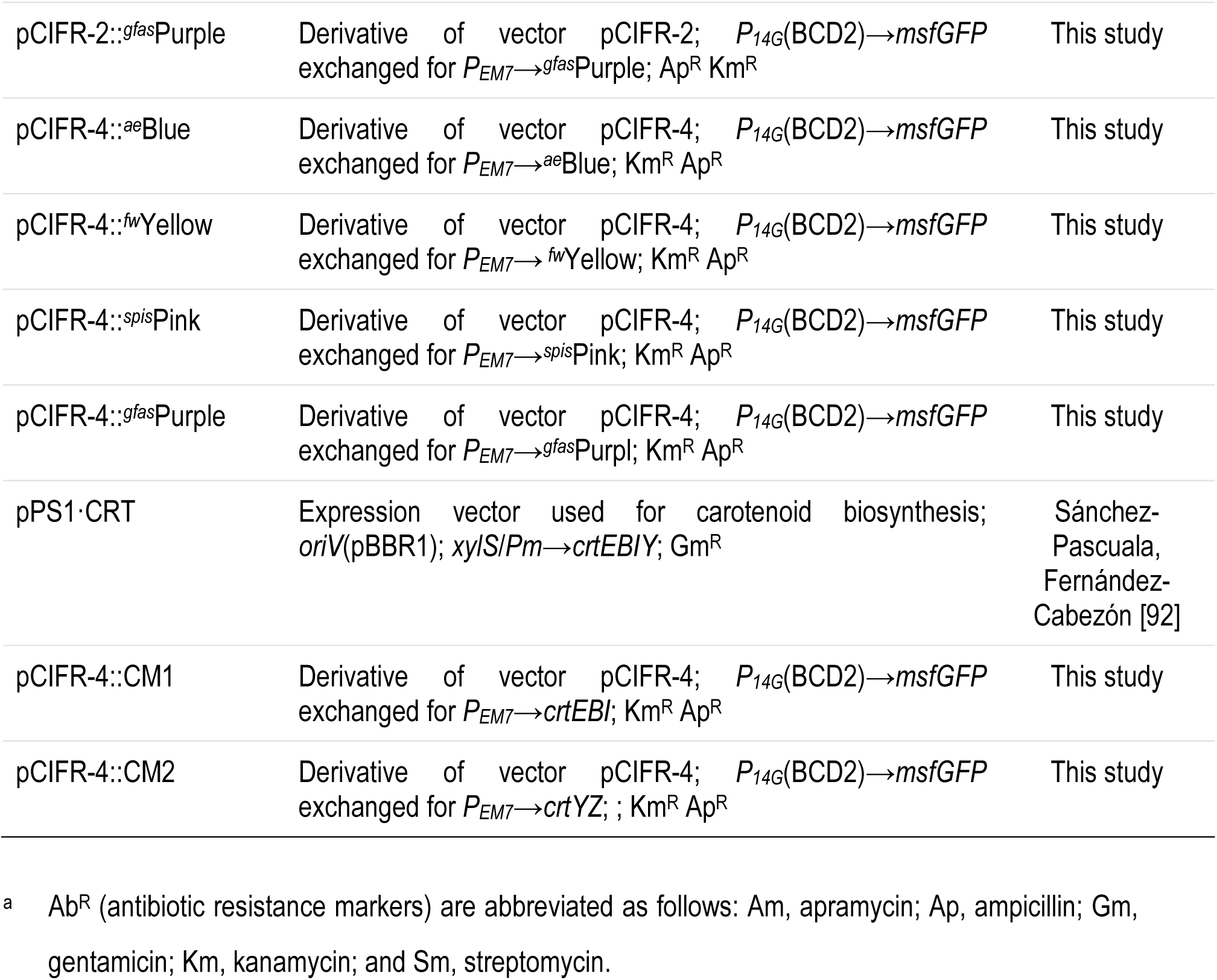
Plasmids used in this study.

Phenotypic characterization in microtiter plate cultures (Epoch 2 plate reader, BioTek Instrument, Winooski, VT, USA) was performed in LB medium; before launching these experiments, cells were pre-grown in the same medium. The precultures were harvested by centrifugation at 8,000×*g* for 5 min and resuspended in the final medium at the starting optical density measured at 600 nm (OD_600_) of 0.01. Cell growth was monitored by measuring the OD_600_ over time [41–45].

### 2.2. Reagents, DNA manipulation, plasmid construction, and DNA sequencing

Chemicals were purchased from Sigma-Aldrich Co. (St. Louis, MO, USA). All oligonucleotides and the genes encoding CrtZ, *^ae^*Blue, *^fw^*Yellow, *^spis^*Pink, and *^gfas^*Purple were synthesized by Integrated DNA Technologies Inc. (Leuven, Belgium). The sequence of all oligonucleotides and synthetic genes is listed in **Table S1** and **S2**, respectively. Synthetic genes were codon optimized for *P. putida* using Benchling (San Francisco, CA, USA). Colony PCR was performed using One*Taq*^TM^ from New England Biolabs Inc. (Ipswich, MA, USA). Uracil-excision (USER) cloning [46] was used for the construction of all plasmids. The AMUSER tool was employed for designing oligonucleotides [47]; Phusion *U* Hot Start^TM^ DNA polymerase from Thermo Fisher Scientific Co. was used according to the manufacturer’s specifications in amplifications intended for USER cloning. The plasmids used in this study are listed in **Table 3**. Plasmid DNA purification was done with the QIAprep Spin Miniprep kit (Qiagen Inc., Valencia, CA, USA) according to the manufacturer’s instructions. DNA Sanger and whole plasmid sequencing were performed at Eurofins Genomics (Ebersberg, Germany) and Unveil Bio (Copenhagen, Denmark), respectively.

**Table 3.** Bacterial strains used in this study.

| Strain | Relevant characteristics<sup>a</sup> | Reference or source |
| --- | --- | --- |
| *Escherichia coli* |  |  |
| DH5α λpir | Cloning host; F<sup>-</sup> λ<sup>-</sup> *endA1 glnX44(AS) thiE1 recA1 relA1 spoT1 gyrA96(Nal<sup>R</sup>) rfbC1 deoR nupG Φ80(lacZΔM15) Δ(argF-lac)U169 hsdR17* (r<sub>K</sub><sup>-</sup> m<sub>K</sub><sup>+</sup>), λpir lysogen | Platt, Drescher [117] |
| ST18 | Helper strain used for conjugation; *pro thi hsdR<sup>+</sup>*; chromosome:RP4-2 Tc<sup>R</sup>::Mu-Kan::Tn7-λpir Δ*hemA*; Sm<sup>R</sup> | Thoma and Schobert [52] |
| HB101/pRK2013 | Helper strain used for conjugation; F<sup>-</sup> λ<sup>-</sup> *hsdS20* (r<sub>B</sub><sup>-</sup> m<sub>B</sub><sup>-</sup>) *recA13 leuB6(Am) araC14 Δ(gpt-proA)62 lacY1 galK2(Oc) xyl-5 mtl-1 thiE1 rpsL20(Sm<sup>R</sup>) glnX44(AS)*; carrying plasmid pRK2013 (*tra<sup>+</sup>* Km<sup>R</sup>) | Figurski and Helinski [53] |
| BW25113 | Wild-type strain; F<sup>-</sup> λ<sup>-</sup> *rrnB3 ΔlacZ4787 hsdR514 Δ(araBAD)567 Δ(rhaBAD)568 rph-1* | Datsenko and Wanner [22] |
| *Pseudomonas putida* |  |  |
| KT2440 | Wild-type strain; derivative of *P. putida* mt-2 [118] cured of the TOL plasmid pWW0 | Bagdasarian, Lurz [119] |
| *Cupriavidus necator* |  |  |
| *C. necator* H16 | Wild-type strain (ATCC 23440) | Schlegel, Kaltwasser [120] |

<sup>a</sup> Ab<sup>R</sup> (antibiotic resistance markers) are abbreviated as follows: Km, kanamycin; Nal, nalidixic acid; Sm, streptomycin; and Tc, tetracycline.

### 2.3. Transformation of E. coli, P. putida, and C. necator

Transformation of chemically-competent or electrocompetent *E. coli* was carried out as described elsewhere [48]. Transformation of *P. putida* was carried out by electroporation [49–51] for replicative plasmids and by conjugation with bi- and tri-parental mating for suicide vectors [18]. Transformation of *C. necator* strains was done by conjugation through bi- and tri-parental mating. Electroporations were performed in a GenePulser XcellTM (Bio-Rad Laboratories, Hercules, CA, USA) configured with a time constant of ∼5 ms and a single pulse of 1.8 kV for *E. coli* (using 1-mm gap width cuvettes) and 2.5 kV for *P. putida* (using 2-mm gap width cuvettes). Biparental mating was carried out using *E*. *coli* ST18 as the donor strain, as described by Thoma and Schobert [52]. Triparental mating was carried out using *E*. *coli* DH5α λ*pir* as the donor strain and *E*. *coli* HB101 carrying plasmid pRK2013 as the helper strain [53].

### 2.4. Mapping the mini-Tn5 transposon insertion sites by arbitrary PCR

The colonies obtained after the insertion of the mini-Tn*5* transposon were used as templates for arbitrary PCR to map the insertion sites within the bacterial genome essentially as described by Martínez-García, Aparicio [54]. The oligonucleotides used for this purpose are listed in **Table S1**. The conditions of the first round of arbitrary PCR were as follows: 5 min at 95°C (initial denaturation); six cycles of 30 s at 95°C, 30 s at 30°C, and 90 s at 72°C; and 30 cycles of 30 s at 95°C, 30 s at 45°C, and 90 s at 72°C [55]. The ARB6 oligonucleotide was used together with the external oligonucleotides within the mini-transposon (*attB*-Ext-F or Sm_Ext_F, for chromoprotein-containing transconjugants with a second insertion with Sm^R^). Then, 1 μL of the first PCR round was used as the template for the second round of arbitrary amplification by applying the following conditions: 1 min at 95°C (initial denaturation); 30 cycles of 30 s at 95°C, 30 s at 52°C, and 90 s at 72°C; followed by an extra extension of 4 min at 72°C. For the second round of arbitrary PCR, the ARB2 oligonucleotide was used together with the internal primer within the mini-transposon (*attB*-Int-F). Finally, the PCR amplification product obtained in the second round was directly sent for Sanger sequencing with the internal oligonucleotide *attB*-Int-F. *P. putida* DNA sequences were analyzed using the *Pseudomonas* Genome Database [56, 57] and BlastN [58]; *E. coli* and *C. necator* DNA sequences were analyzed using BlastN. Additional information on the genes found was obtained with KEGG [Kyoto Encyclopedia of Genes and Genomes] [59] and BioCyc [60].

### 2.5. Plate reader experiments and fluorescence quantification

Before launching plate reader experiments, cells were pre-grown in 1 mL LB medium in 96 deep-well plates. After measuring the OD_600_, the precultures were harvested by centrifugation at 8,000×*g* for 5 min and resuspended in LB at the starting OD_600_ of 0.01. The cells were then transferred to 96-well microtiter plates and incubated at 30°C in a plate reader (Synergy H1, BioTek Instruments). The fluorescence of msfGFP was monitored using an excitation wavelength (λ_excitation_) of 488 nm and an emission wavelength (λ_emission_) of 510 nm. The fluorescence of *^fw^*Yellow was monitored at λ_excitation_ = 520 nm and λ_emission_ = 540 nm. To normalize fluorescence data, cell growth was also monitored by measuring the OD_600_ over time; the fluorescence of cultures of KT2440 was used as blank for each time point.

### 2.6. Extraction and analysis of carotenoids

Selected transconjugants were grown in 1 mL LB medium, in 96 deep-well plates. Carotenoids were extracted from cell pellets as previously reported [61]. After measuring the OD_600_, the cultures were harvested by centrifugation at 4,500×*g* for 15 min. The cell pellet was washed with water, centrifuged again at 4,500×*g* for 15 min and then extracted in 500 μL of acetone at 55°C for 30 min under agitation, covered in aluminum foil to prevent light degradation. The mixture was then centrifuged at 11,000×*g* for 1 min, and the supernatant was transferred into another microcentrifuge tube and stored at –20°C for analysis [62]. To analyze the lycopene (Lyc) content of the Lyc-producing strains, 200 μL of the acetone supernatants were transferred to a 96-well microtiter plates and their absorbance at 474 nm was measured and compared to the values obtained with a standard curve. To derive the carotenoid content per cell, the value obtained was divided by the OD_600_ of the corresponding culture. To analyze the total carotenoid content, the acetone extracts were dried using an Eppendorf Concentrator Plus (Eppendorf, Hamburg, Germany) at 45°C and 1,250×*g* for 3 h. The samples were resuspended in 1 mL of 99% (v/v) ethanol and transferred to glass vials. Calibration curves for carotenoid quantification were prepared from commercial standards of Lyc, β-carotene, and zeaxanthin (Zea). The calibration curves were prepared in 99% (v/v) ethanol (1 μg mL^−1^, 5 μg mL^−1^, 10 μg mL^−1^, 25 μg mL^−1^, 50 μg mL^−1^, and 100 μg mL^−1^). The calibration curve samples were then analyzed alongside of the biological samples using high performance liquid chromatography (HPLC). The HPLC-system (Ultimate 3000, Thermo Fisher Scientific, Waltham, MA, USA) was equipped with a Supelco Discovery^TM^ HS F5 column (2.1×150 mm, particle size = 3 μm) and the chromatographical separation was performed using 10 mM ammonium formate (pH = 3.0, adjusted with formic acid) as eluent A and acetonitrile as eluent B. The ammonium formate and the formic acid were both LC-MS grade and they were purchased from Sigma Aldrich (Merck Life Science A/S, Søborg, Denmark) while the LC-MS grade acetonitrile was purchased form VWR (Avantor, Radnor, PA, USA). Gradient elution was applied at a flow rate of 0.7 mL min^−1^ according to the following: 0-2 min 25% B, 2-4 min 25% to 90% B, 4-10.5 min 90% B, and 10.5-11.7 min 90% to 25% B. The column was then re-equilibrated at 25% B for 1.8 min. The injection volume for each sample was 10 μL and the column oven was maintained at 30°C. The detection of the carotenoids was done using a DAD-300 detector at a λ = 450 nm. All data was acquired and processed using the Chromeleon^TM^ software (version 7.2.9; Thermo Fisher Scientific). The content of each carotenoid in the samples was normalized by dividing the acquired quantified concentrations by the estimated cell dry weight (CDW). The CDW was obtained from the OD_600_ using a conversion factor of 0.533 [63, 64].

## 3. RESULTS AND DISCUSSION

### 3.1. Design and assembly of the CIFR system

Transposon tools have been extensively developed and utilized for metabolic engineering. A significant limitation of existing Tn*5*-based systems is the inability to recycle functional components for subsequent rounds of insertions. To address this issue, we designed the CIFR system for iterative bacterial engineering, comprising two sets of plasmids: integration vectors (pCIFRs) and helper plasmids (pFNCs) (**Fig. 1a**). The pCIFR integration vectors (**Fig. 1b** and **Fig. S1a**) are derived from the pBAM (*borne-again mini-transposon*) plasmids [65] and their pBAMD-derivative vectors [54]. The Ab^R^ cassettes in the transposon module are equipped with *attP*/*attB* sites flanking the resistance genes, facilitating clean excision. Sharing structural and functional elements with pBAMD1s, pCIFR vectors include the following features: (i) an ampicillin resistance (Ap^R^) cassette outside the transposon module for cloning and counter-selection of plasmid replication; (ii) the narrow host-range origin of replication of plasmid R6K (*ori*R6K), which requires a *pir*^+^ or *pir116* strain for replication (Π protein dependent); (iii) the origin of transfer *oriT*, allowing for the conjugative transfer of the plasmid from the host (cloning) strain to a new bacterial recipient through *tra*-mediated mobilization; (iv) the hyper-active Tn*5* transposase gene (*tnpA*) [66]; (v) a mini-transposon module, delimited by two mosaic elements (termed ME-I and ME-O to distinguish their orientation but otherwise identical in sequence), containing the multiple cloning site (MCS), a monomeric superfolder GFP gene (*msfGFP*), expressed under the control of the strong regulatory elements *P_14G_* and BCD2 [67, 68], and an Ab^R^ cassette (X^R^) used as a marker for the integration event, flanked by the *attP* and *attB* sequences. The pCIFR plasmids provide four selection markers conferring resistance to Km, Sm, Gm, or Am. These plasmids are termed pCIFR-2, pCIFR-4, pCIFR-6, and pCIFR-8, respectively, using the same code for the Ab^R^ determinants as in the Standard European Vector Architecture [69, 70]. Throughout this work, we will collectively refer to the four plasmid variants as pCIFR-*X* (**Fig. 1a**).

**Fig. 1.**
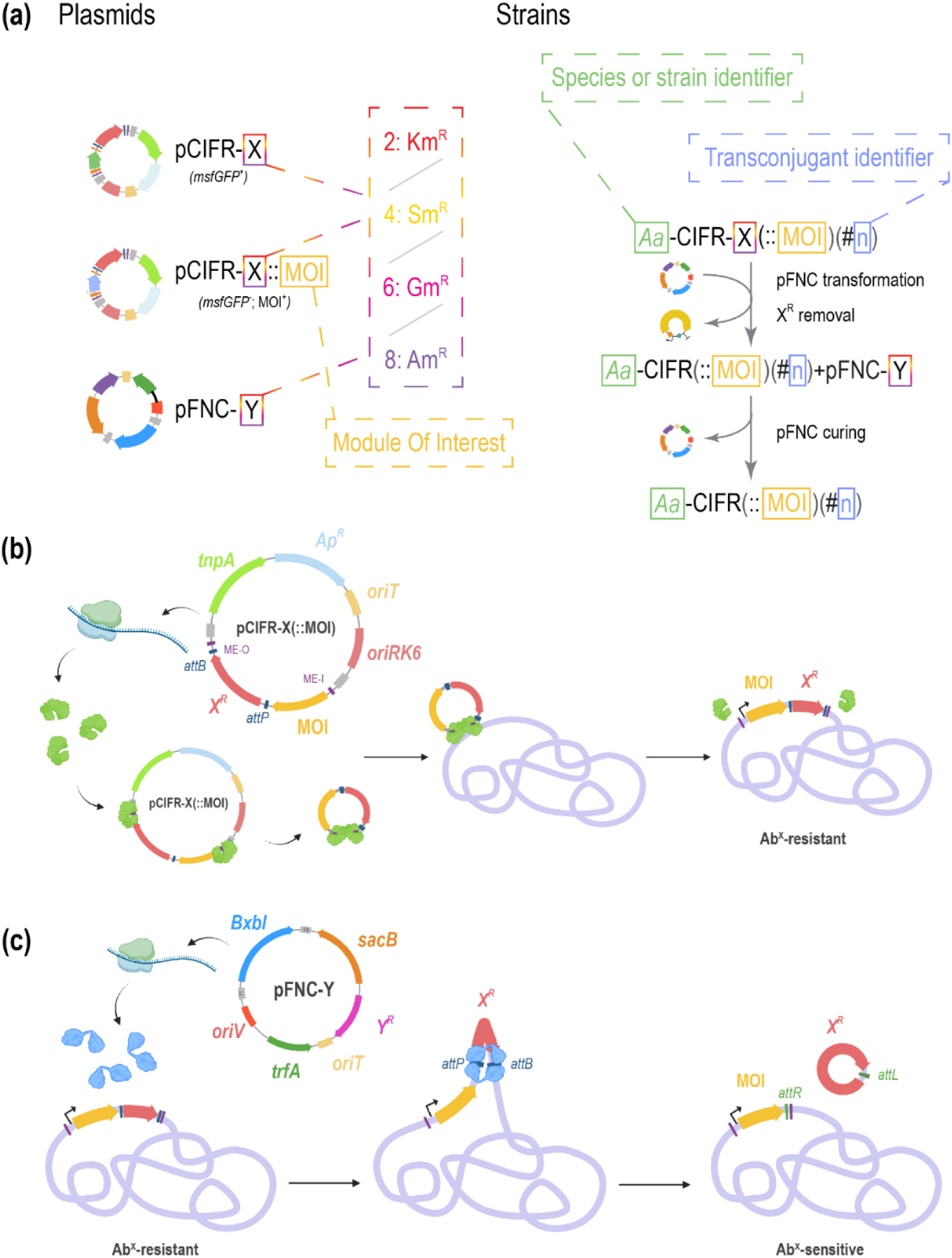
Nomenclature and functioning of pCIFRs and pFNCs plasmids. **(a)** Nomenclature scheme for the different plasmids (left) and strains (right) presented in this study. In pCIFR-X and pFNC-Y, X and Y represent the antibiotic resistance (Ab^R^) cassette that is available for the plasmids. The choice is between 2, KmR, red; 4, SmR, yellow; 6, GmR, pink and 8, AmR, purple. When a module of interest (MOI) gets cloned into a pCIFR-X, a pCIFR-X::MOI is generated. A strain engineered with a pCIFR-X is a “Aa-CIFR-X”, where Aa stands for an identifier for the species or strain. If the pCIFR-X contains a MOI instead of a *msfGFP* expression cassette, “::MOI” is added to the name of the strain. After a transformation with pCIFR-X(::MOI), every transconjugant is genetically distinct from the others, and it can be identified with a different number (#n). When a pFNC-Y is transformed into one of these TJs to flip out the X^R^ cassette, “-X” is removed from “p-CIFR-X” and “+pFNC-Y” is added to the name of the strain. Upon sucrose-mediated curing of pFNC, “+pFNC-Y” is removed from the name, leaving “Aa-CIFR(::MOI)(#n)”. The name parts in a box change depending on the case and the parts between brackets are optional. **(b)** Representation of the insertion of the Tn*5* mini-transposon in the bacterial genome performed by the TnpA transposase. Upon transient expression of *tnpA* in the target organism, the transposase recognizes the mosaic elements and, while still binding the DNA, dimerizes, forming the synaptic complex. The transposon sequence is then cleaved and inserted in a “random” site in the chromosome(s) of the host. The cells in which the transposition has been successful can be selected thanks to the selection marker X^R^ present in the transposon module. **(c)** Representation of the flip-out of the Ab^R^ cassette from the genome performed by the Bxb1 phage integrase. The integrase can recognize the sites *attP* and *attB* and perform a recombination between them, contextually excising from the genome the sequence contained in between (in this case, X^R^). Since the excised sequence does not contain an origin of replication, the Ab^R^ phenotype is lost as the cells duplicate. The recombination between *attP* and *attB* creates the hybrid sites *attR* and *attL*, that cannot be recognized by the integrase in absence of a directionality factor. In this way, the system can be used repetitively without risking a cross recombination between *att* sites. Panels **(b)** and **(c)** were created with BioRender.com.

The excision of the transposon-encoded resistance cassette is facilitated by a second set of plasmids, named pFNCs (“Flip aNd Cure”; **Fig. 1c** and **Fig. S1b**). The functionality of these plasmids relies on two key features: the gene encoding Bxb1-int, which catalyzes recombination between the *attP* and *attB* motifs, and the *sacB* gene from *Bacillus subtilis*, which encodes levansucrase. Levansucrase hydrolyzes sucrose to produce levan polysaccharides, rendering the cells sensitive to sucrose due to the toxicity of the levan polymers [71]. The pFNC-*Y* plasmids are constructed on a pSEVA*Y*213 backbone, where *Y* is a placeholder representing one of four possible Ab^R^ cassettes (Km, 2; Sm, 4; Gm, 6; and Am, 8; as indicated for pCIFR plasmids, **Fig. 1a**). These plasmids include the RK2 replication module, consisting of the origin of vegetative replication (*oriV*) and *trfA*, encoding the TrfA protein that activates *oriV* [72]. Additionally, the pFNC-Y vectors contain an origin of transfer (*oriT*), allowing for conjugative mobilization. The *sacB* gene, enabling the selection of successful excision events, is positioned in the gadget position of the SEVA architecture to confer sucrose sensitivity.

### 3.2. A four-step protocol for iterative genome engineering of Gram-negative bacteria

After cloning and confirming the identity of all plasmids in the toolset, we established a streamlined protocol for their use, captured by the acronym CIFR—*Clone*, *Integrate*, *Flip-out*, and *Repeat*. The protocol can be divided into four operative parts (**Fig. 2**).

**Fig. 2.**
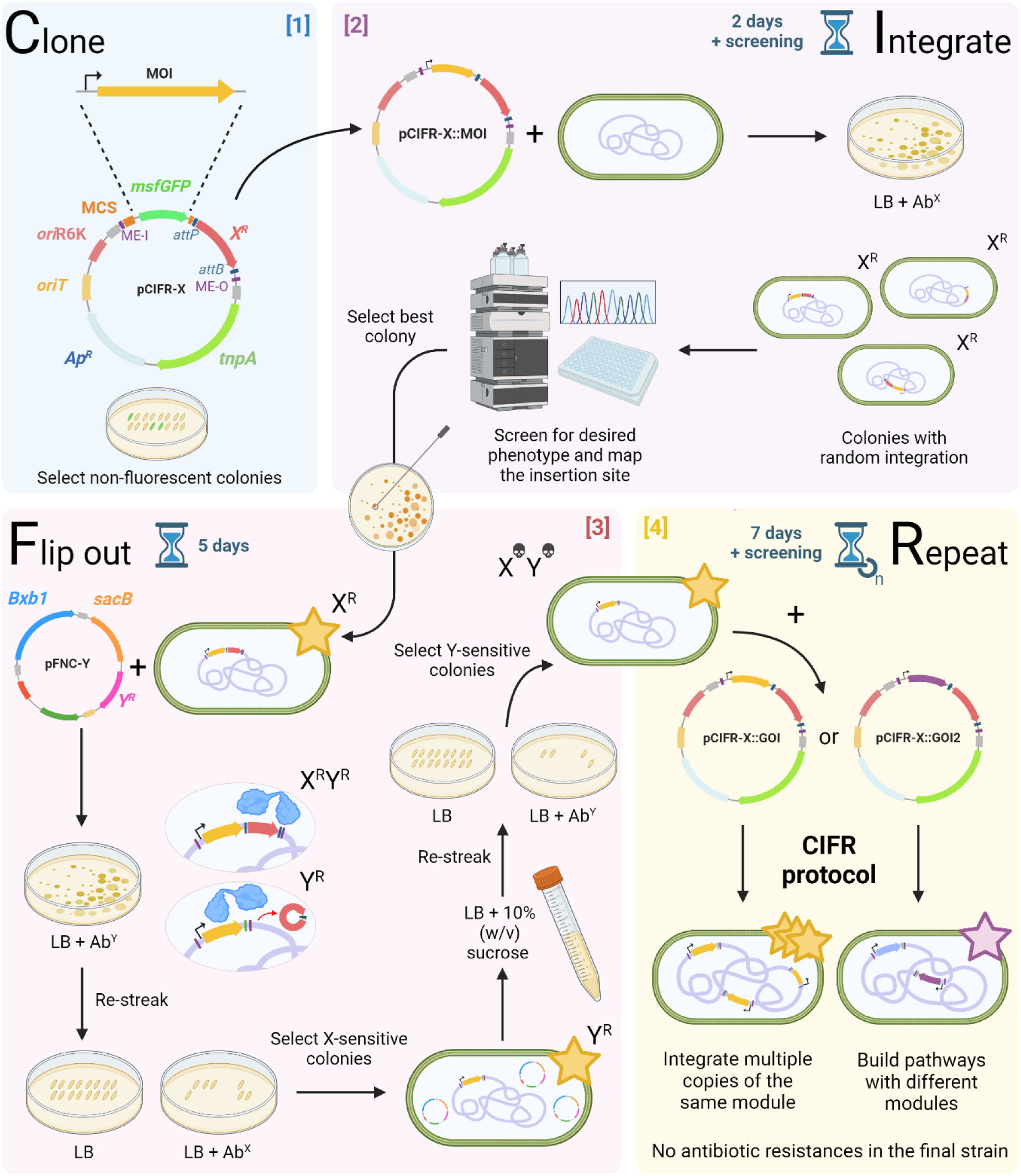
CIFR facilitates genome engineering of Gram-negative bacteria in an iterative fashion through a four-step [1–4] protocol. [1] ***CLONE*:** The pCIFR-X suicide vector harbors a synthetic Tn*5*-transposase gene (*tnpA*) and a transposon module flanked by mosaic elements, which can integrate into the genome of various Gram-negative bacteria. The transposon module includes one of four antibiotic resistance markers (kanamycin, Km; streptomycin, Sm; gentamicin, Gm; and apramycin, Am) for selecting integrants, and an *msfGFP* gene constitutively expressed under the strong P_14G_(BCD2) element. This fluorescent marker facilitates the identification of non-fluorescent colonies in the cloning host. The marker can be swapped for the module of interest (MOI, that can be a gene or an operon). [2] *INTEGRATE*: The pCIFR-X::MOI plasmid is transformed into the host *via* electroporation or conjugation. As the plasmid can only replicate in λ*pir* strains, antibiotic-resistant colonies of the target host carry the integrated transposon module in their chromosome(s). Desired phenotypes can be screened at this stage, and transconjugants selected for further characterization. Integration sites can be mapped using an arbitrary PCR protocol with degenerate primers. [3] *FLIP OUT*: To facilitate further engineering, the resistance marker can be excised from the chromosome by transforming selected transconjugants with pFNC-Y. This plasmid encodes the BxbI phage recombinase, which recognizes *attP* and *attB* sites flanking the X^R^ cassette and mediates recombination. X-sensitive colonies are selected by replica plating on media with and without antibiotic X. pFNC-Y can carry different antibiotic resistances, and includes a counterselection marker, *sacB*, for plasmid removal by overnight cultivation with 10% (w/v) sucrose. Colonies that lost the plasmid are identified by replica plating on media with and without antibiotic Y. [4] *REPEAT*: Since the X^R^ is removed and the *att* sites are altered upon the first recombination, the protocol can be iterated to increase the copy number of integrated module(s) or integrate separate modules of a complex pathway. A single iteration of the CIFR protocol takes 7 days. ME, mosaic element; MCS, multiple cloning site; MOI, module of interest; X^R^, resistance cassette for antibiotic X (pCIFR plasmids); and Y^R^, resistance cassette for antibiotic Y (pFNC plasmids). Figure created with BioRender.com.

#### [1] Clone

a) The cloning step in pCIFR vectors is facilitated by the various restriction sites present in the MCS and at the 3′-end of *msfGFP*, inherited from pBAMD plasmids. These sites allow the insertion of the desired construct between the ME-I and the *attP* sequences, thereby replacing *msfGFP*. Cloning can also be performed using restriction enzyme-independent methods, such as Gibson assembly or USER cloning [73, 74]. After the cloning steps, pCIFR-X becomes pCIFR-X::MOI (module of interest, **Fig.1a**).
b) The presence of *msfGFP* aids in screening for the desired colonies during the cloning process. msfGFP production is enhanced by the strong *P_14G_* promoter and BCD2, making fluorescent colonies easy to identify [75, 76]. Clones that do not exhibit green fluorescence under blue light are those where *msfGFP* has been replaced with the gene or operon of interest.

#### [2] Integrate

a) After selecting the non-fluorescent clones of *E. coli* DH5α and confirming the sequence of the inserted gene, the next step is to integrate the gene of interest into the target genome. This part of the protocol mirrors the integration process used with pBAMD vectors. The transformation of pCIFR plasmids into the target organism can be accomplished by various methods, with conjugation being the most effective. Electroporation is also possible but less efficient, as pCIFRs cannot replicate in strains lacking the gene encoding the Π protein. Upon transformation, the transient expression of *tnpA* facilitates the transfer of the transposon module to the target genome (**Fig. 1b**). The integrated segment between the two mosaic elements includes the MOI and X^R^, i.e. the resistance cassette to antibiotic X (Ab^X^, as in pCIFR-X, following the SEVA nomenclature). Plating or streaking on LB plates containing Ab^X^ (at the appropriate concentration) is used to select for successful integration events. If pCIFR-X is transformed via conjugation, it is also necessary to select against the donor strain (and helper strain in the case of triparental conjugation; see *Materials and Methods*).
b) A screening procedure is essential for selecting transconjugants (TCJs) with the desired phenotypic traits after obtaining a population with different integration locations. This step is the most time-consuming, as it depends on the analytical techniques employed. To expedite the process, we recommend using fluorescent or colored markers, or growth-coupled selection if the phenotype is suitable [77]. During the screening, the population remains resistant to Ab^X^, which helps prevent contamination. At the end of this screening procedure, the desired TCJ(s) should be isolated and used for subsequent steps.

#### [3] Flip-out

a) The flip-out procedure allows for the removal of the Ab^R^ gene from the target genome in two steps. The first step involves transforming the selected TCJ(s) with one of the pFNC plasmids. The selected pFNC-Y plasmid should contain an Ab^R^ cassette (Y^R^) different from the one used in the pCIFR-X integration step. Additionally, both X^R^ and Y^R^ markers should differ from any Ab^R^ naturally present in the host strain or used by other plasmids or genomic integrations. Both electroporation and conjugation are possible for this transformation step, thanks to the presence of *oriT* in the backbone of pFNC vectors. In this case, TCJs are selected on LB plates supplemented with antibiotic Y (Ab^Y^). The use of pCIFR-6/8 with pFNC-6/8 is not recommended, since the Am^R^ cassette (8) also confers resistance against Gm [78].
b) The *bxb1-int* gene is expressed in transformants harboring pFNC-Y, producing Bxb1-int, which recognizes the *attP* and *attB* sites, performs a recombination on them and excises the *X*^R^ cassette (**Fig. 1c**). Since the excised sequence lacks an origin of replication, the resulting small circular DNA cannot be propagated by the cells, leading to the loss of Ab^R^ during cell growth. To select transformants where the flip-out has occurred, they are re-streaked onto LB plates and LB plates containing Ab^X^. Clones sensitive to Ab^X^ are those where the *X*^R^ has been successfully removed from the bacterial genome. The recombination between *attP* and *attB* yields an *attR* scar, that cannot be recognized by Bxb1-int in absence of a directionality factor [79].
c) The final step involves curing the pFNC-Y plasmid. This process is facilitated by the presence of the counter-selection marker *sacB*, which is used to select against cells harboring the plasmid in a culture medium with a high sucrose concentration. One of the Ab^X^-sensitive clones is inoculated in LB medium with 10% (w/v) sucrose and incubated overnight. From this culture, cells are streaked on an LB plate and clones are screened for Ab^Y^ sensitivity, similarly to the previous step. The obtained Ab^Y^-sensitive clones are those cured of the pFNC-Y plasmid.

#### [4] Repeat

a) This final step is optional, as the strain obtained after the flip-out may already exhibit the desired phenotype. However, the ability of the CIFR system to perform multiple integration events, even with the same plasmid, can be exploited for further engineering steps. The final strain produced *via* the CIFR protocol is free of any Ab^R^ or *attP* and *attB* sites, allowing for further iterations with either pCIFR-X or other plasmids using the same Ab^R^ marker. This flexibility enables the construction of complex phenotypes through the random integration of different functional modules, since each iteration results in varying expression levels due to the combination of user-designed regulatory elements and integration loci. Additionally, the same module can be integrated multiple times, enhancing the phenotype associated with its expression.

### 3.3. A uniform nomenclature for strains constructed with the CIFR protocol

To avoid confusion with the different steps of the CIFR protocol, the following nomenclature was introduced to refer to the different strains (**Fig. 1a**):

- *(Name of the species or strain)*-CIFR-X is an engineered strain with a random integration using pCIFR-X, bearing the reporter gene *msfGFP* and X^R^ (after step [2] of the protocol).
- *(Name of the species or strain)*-CIFR+pFNC-Y is an engineered strain with a random integration performed with pCIFR-X, containing only the reporter gene *msfGFP* and X^R^ flipped-out with pFNC-Y (during step [3] of the protocol). After the curing of the helper plasmid, “+pFNC-Y” is removed from the name of the strain (after completing step [3] of the protocol).
- *(Name of the species or strain)-*CIFR-X::MOI is an engineered strain with a random integration performed with plasmid pCIFR-X::MOI, bearing the module of interest and X^R^.
- *(Name of the species or strain)*-CIFR::MOI+pFNC-Y is an engineered strain with a random integration with plasmid pCIFR-X::MOI, bearing the module of interest, but with X^R^ flipped-out with pFNC-Y (during step [3] of the protocol). After the curing of the helper plasmid, “+pFNC-Y” is removed from the name of the strain (after completing step [3] of the protocol.
- During and after the phase of selection and screening (end of step [2]), the different TCJs can be identified with a # followed by a progressive number.

### 3.4. Heterologous DNA integration in the genome of three Gram-negative bacteria using the CIFR toolset

The CIFR system was tested in three different Gram-negative hosts commonly used in metabolic engineering: the soil bacterium *P. putida* KT2440, the laboratory workhorse *E. coli* BW25113, and the chemoautotroph *C. necator* H16. Since applying the plasmid toolset requires prior knowledge of the Ab^R^ profile of the host organism, we conducted an antibiotic sensitivity test for *C*. *necator*. Besides Km, Cm, and Tc, previously used as antibiotic markers as reported in the literature [80, 81], other Ab^R^ were unknown. We cultivated *C*. *necator* on LB agar plates and in 96-well microtiter plates with varying concentrations of commonly used antibiotics (**Fig. S2** and **Table S3**). The results indicated that Km at 100 μg mL^−1^, Cm at 50 μg mL^−1^, Sm at 200 μg mL^−1^, and Tc at 15 μg mL^−1^ can be used for selection in *C*. *necator* H16. Ampicillin, Gm, Am, and 5-chloro-2-(2,4-dichlorophenoxy)phenol (Irgasan) were ineffective at inhibiting microbial growth, even at the highest concentrations tested (400, 40, 100, and 50 μg mL^−1^, respectively).

We then transferred a set of pCIFR vectors to the target bacterial host *via* conjugation. For *P. putida*, triparental conjugation was performed using *E. coli* HB101 carrying plasmid pRK2013 as the helper strain (**Tables 2** and **3**). For *E. coli* BW25113 and *C. necator*, biparental conjugation was used, with *E. coli* ST18 serving as the helper strain [52, 53]. All pCIFR variants were tested in *P. putida*, while *E. coli* and *C. necator* were transformed with vector pCIFR-2. After counterselecting for the donor and helper strains on appropriate culture media (see *Materials and Methods*), fluorescent colonies of the recipient strains could be easily identified, confirming successful conjugation and transposition. This result also indicated that the addition of the *attP*/*attB* sites flanking the mini-Tn*5* module did not interfere with the transposase ability to mediate the integration of the transposon DNA into the bacterial genome.

We isolated 64 independent TCJs of *P. putida*-CIFR-X (termed *Pp*-CIFR-X), selecting 16 TCJs for each CIFR variant. For *E. coli* and *C. necator*, 48 TCJs were isolated and retained for further analysis (named *Ec*-CIFR-2 #n and *Cn*-CIFR-2 #n, respectively). Arbitrary PCR was used to map the positions of the different transposon integrations *via* Sanger sequencing as described by Das, Noe [55]. We observed certain hotspots where integrations were more frequent (**Tables S5-S7** and **Fig. 3a**). A common hotspot for all three organisms was the region located near the origin of replication of the chromosome(s), with more than 75% of the total insertion sites found closer to the origin than to the terminus. Except for the pHG1 megaplasmid of *C. necator*, approximately 60% of all mini-Tn*5* insertions occurred within 1 Mb of the origin of replication. This phenomenon is likely due to this part of the chromosome being present in multiple copies in growing cells. Since this segment is involved in replication, it exists in a more open structure than the rest of the chromosome, making it more accessible for the transposase to perform integration [82, 83].

**Fig. 3.**
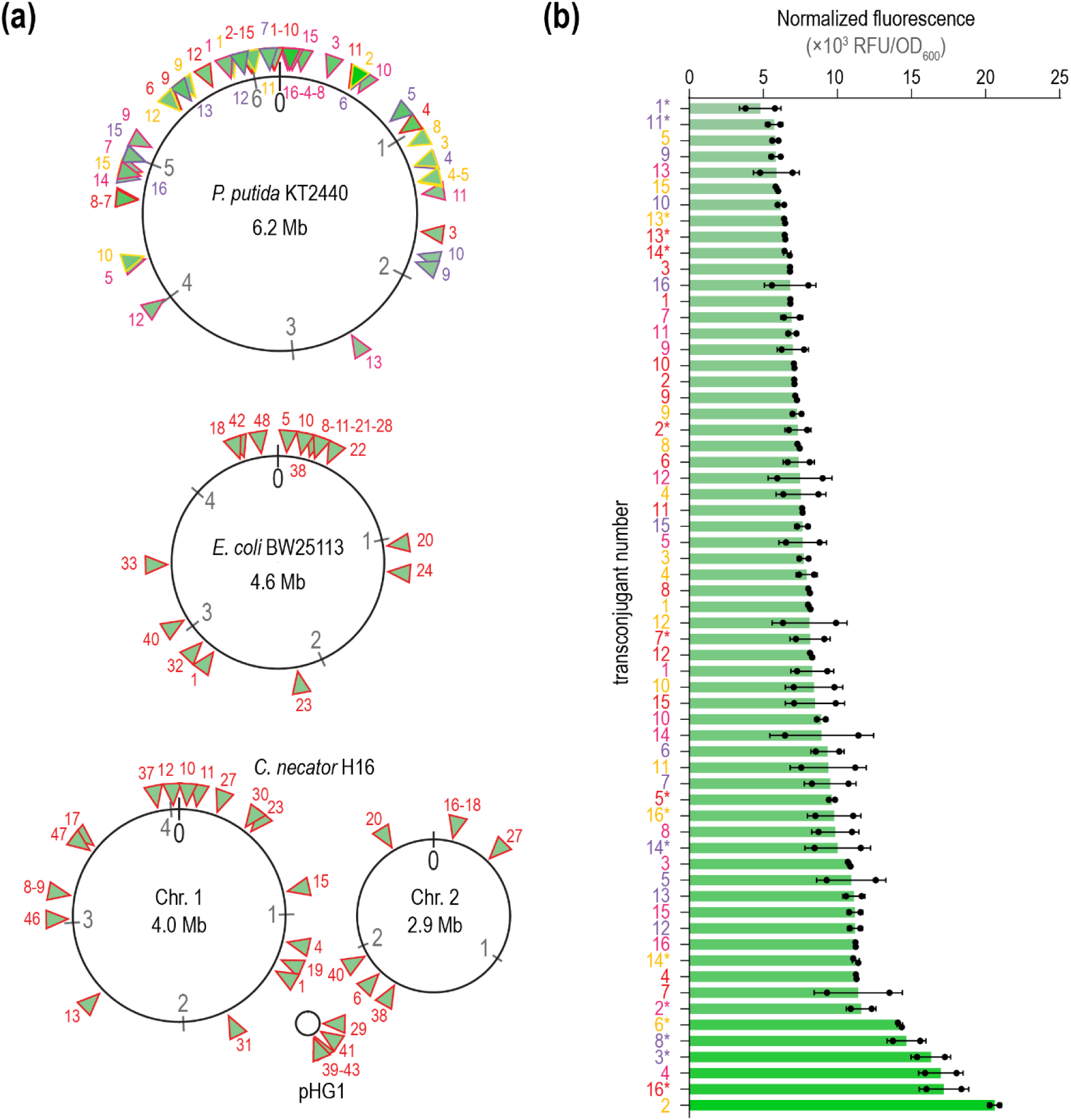
Characterization of mini-Tn5 integration in three different Gram-negative hosts. **(a)** The integration of *msfGFP* using CIFR in *P. putida* KT2440, *E. coli* BW25113, and *C. necator* H16 was mapped via arbitrary PCR. The positions of individual integrations in the chromosomes of the different hosts are indicated as triangles. Each triangle is numbered to identify transconjugants (TCJs) obtained with different pCIFR plasmids. The color for the border and label of each triangle denotes the antibiotic used for integration. For *P. putida*, the green hue of each triangle is proportional to the relative fluorescence intensity exhibited by the TCJ carrying that integration. (b) Relative *msfGFP* fluorescence, normalized to the optical density at 600 nm (OD_600_), of individual *P. putida* KT2440 TCJs. Each column label identifies TCJs obtained via transposition with different pCIFRs, with colors representing the antibiotics used for integration as follows: red: CIFR-2 (Km); yellow: CIFR-4 (Sm); purple: CIFR-6 (Gm); and violet: CIFR-8 (Am). Data represent average values from 2 independent measurements ± standard deviation.

We further characterized the effect of different insertion sites on reporter gene expression in *P. putida* TCJs, testing all CIFR plasmids. Each vector contains the *P_14G_*(BCD2)→*msfGFP* module for strong expression of the reporter gene. The fluorescence of the 64 isolated colonies was analyzed in liquid LB cultures using a microtiter plate reader. Background fluorescence from strain KT2440 was used as a blank, and fluorescence readings (λ_excitation_ = 488 nm/λ_emission_ = 510 nm) were normalized to the OD_600_ of the cultures (**Fig. 3b**). Under these conditions, we observed up to a 4-fold difference between the lowest and highest maximum normalized fluorescence values, with most colonies displaying a range of roughly a 2-fold variation in msfGFP production. TCJ #2 from the *Pp*-CIFR-4 TCJ population exhibited the highest relative fluorescence, with the integration occurring within the *PP_0411* gene, which encodes an ATP-binding protein of a polyamine ABC transporter. Overall, we found no clear correlation between fluorescence intensity and chromosomal position (**Fig. S3**). Taken together, these results show that the CIFR system can be used to integrate expression cassettes into the genomes of three different Gram-negative bacteria in a random fashion.

### 3.5. The helper pFNC plasmids mediate the excision of antibiotic resistance cassettes

We conducted experiments to validate the efficiency of the flip-out step, a unique feature of the CIFR protocol, using the TCJs described in the previous section that carry a *msfGFP* expression cassette in their genome. In each case, 48 TCJs representing different combinations of target organisms and pCIFR variants (16 for *Pp-CIFR-2*, 16 for *Pp-CIFR-4*, 16 for *Ec-CIFR-2*, and 16 for *Cn-CIFR-2*) were selected. Each group of 16 TCJs was further divided into three smaller groups of 5-6 TCJs, pooled together, and inoculated in LB supplemented with the appropriate antibiotic (Km or Sm) to prepare pre-cultures for transformation with the helper pFNC-Y plasmid. *P. putida* and *E. coli* were transformed *via* electroporation; *C. necator* was transformed *via* conjugation. Pooling TCJs helped avoid potential bias from variations in flip-out efficiency due to the genomic location of the integration. To select for transformed cells, the mixtures were plated on LB plates supplemented with the appropriate antibiotic to select for pFNC-Y (Ab^Y^). Next, 50 single colonies from each plate were randomly selected and re-streaked on both LB and LB + Ab^X^ (Km/Sm) plates, depending on the Ab^R^ cassette encoded on the corresponding pCIFR vector. Results for *Pp*-CIFR-2/4+pFNC-6/8 are shown in **Fig. 4a** as an example. Transformants that could grow on the antibiotic plates still harbored the Ab^R^ cassette along with the *msfGFP* module. In contrast, transformants sensitive to the antibiotics were those in which Bxb1-int successfully mediate the flip-out.

**Fig. 4:**
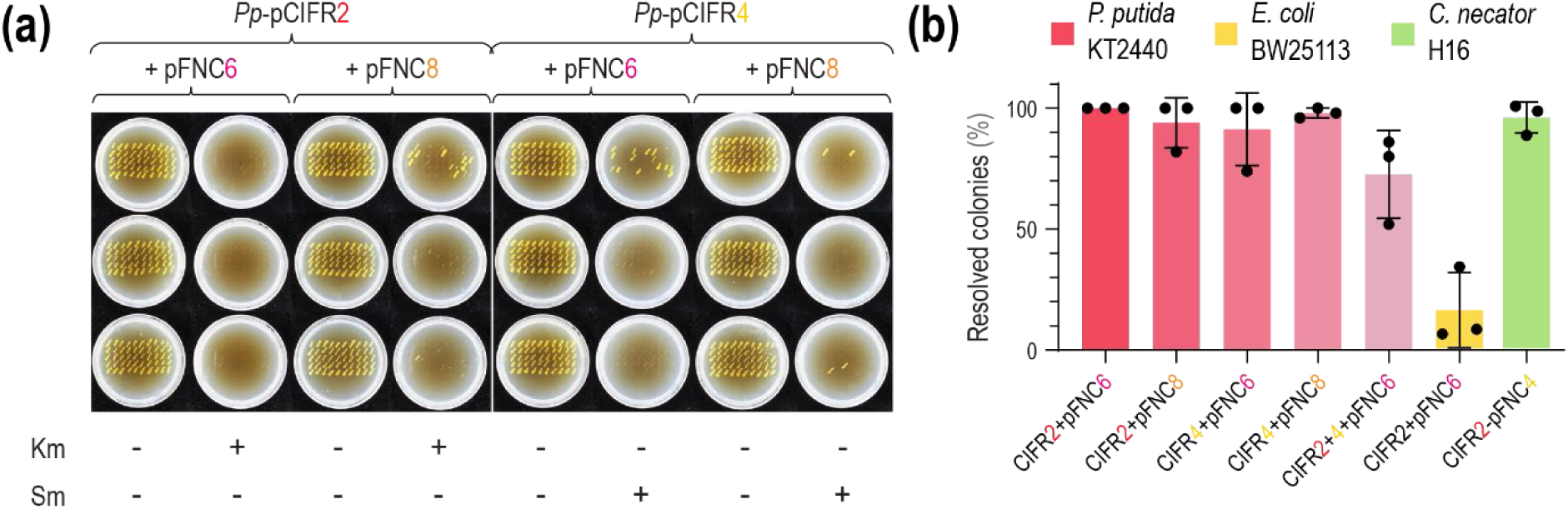
Efficiency of antibiotic cassette removal by pFNC-mediated excision. (a) After transforming *P. putida* KT2440 with plasmid pCIFR-2/4, three pools of 5-7 colonies were inoculated and transformed with plasmid pFNC6/8. Single colonies were isolated, and 50 clones from each transformation were replica-plated on LB plates with and without antibiotics (Km for plasmid pCIFR-2, Sm for plasmid pCIFR-4). The efficiency of antibiotic cassette removal mediated by pFNC-Y was determined by dividing the number of antibiotic-sensitive clones by the total number of colonies tested. This operation was performed for each combination shown in panel **(b)**, with one representative set of plates displayed as an example. **(b)** Resolution efficiency of the different combinations of pCIFR-X and pFNC-Y plasmids in the different bacterial hosts. The color code for plasmid numbers, referring to the different plasmid variants, is the same is indicated in Fig. 2. Plasmid pFNC could resolve two antibiotic resistances simultaneously in *P. putida* KT2440 transconjugants, albeit at a lower efficiency. Data represent average values from three independent measurements ± standard deviation.

The flip-out efficiency was calculated as the percentage of Ab^X^-sensitive transformants out of the 50 that were tested. The efficiencies for all combinations are shown in **Fig. 4b**. In *P. putida* and *C. necator*, the Bxb1-int achieved the removal of the Ab^R^ cassette with efficiencies of 96 ± 8% and 95 ± 6%, respectively. In *P. putida*, different combinations of pCIFR-X/pFNC-Y were tested. For two of these combinations, the efficiency was slightly lower in one out of three replicates (**Fig. 4a**), bringing the average efficiency to 94 ± 10% for *Pp*-CIFR-2+pFNC-8 and 91 ± 15% for *Pp*-CIFR-4+pFNC-6. Other combinations showed even higher efficiencies, with 100 ± 1% for *Pp*-CIFR-2+pFNC-6 and 98 ± 2% for *Pp*-CIFR-4+pFNC-8. In *E. coli*, the flip-out process occurred at a lower efficiency (16 ± 5%), which may be related to the copy number of pFNC plasmids in the different organisms. These replicons are reported to be maintained at ∼30-40 copies per genome in *P. putida* and *C. necator* [84], but this value is an order of magnitude lower in *E. coli* [85, 86]. Additionally, other regulatory elements, such as the promoter and ribosome binding site (RBS), could influence protein abundance and the effectiveness of the integrase in different organisms. In any case, at the flip-out stage of the CIFR protocol, the desired phenotype should already have been selected. Hence, achieving a high efficiency is not critical—obtaining even one antibiotic-sensitive clone to continue with the protocol may be sufficient depending on the application. We recommend screening at least 50 colonies to verify flip-out if the target organism has a low pFNC copy number. A smaller number of colonies may suffice when pFNC shows a medium or high copy number, as in the cases of *C. necator* and *P. putida*.

Next, we evaluated the ability of pFNC to flip out two Ab^R^ cassettes simultaneously. This feature could streamline the CIFR protocol by uncovering the desired phenotype resulting from the combination of two different modules randomly inserted into the genome. Multiple integrations using a Tn*5* mini-transposon are feasible while selecting for different Ab^R^s, as the transposase typically acts in *cis* and does not further process MEs already present in the genome [65]. To test this scenario, a random TCJ from the *Pp*-CIFR-2 group was selected and transformed with plasmid pCIFR-4 through triparental conjugation. Sixteen colonies of *Pp*-CIFR-2+CIFR-4 were then randomly selected, pooled into three groups of 5-6 colonies, and used to inoculate precultures for transformation with the helper plasmid pFNC-8. Fifty colonies from each group of *Pp*-CIFR-2+CIFR-4+FNC8 were then re-streaked on LB, LB containing Km, and LB containing Sm. An illustrative example of the results is shown in **Fig. S4**. For calculating the flip-out efficiency of the helper pFNC plasmids, only transformants sensitive to both antibiotics were considered.

While the efficiency was lower than when removing a single Ab^R^ cassette, the process was still highly efficient with an average of 73 ± 18% sensitive colonies (**Fig. 4b**).

We then evaluated the precision of Bxb1-int during the flip-out step by examining the scar left by the CIFR system in the bacterial genome. Theoretically, after the removal of the Ab^R^ cassette, the DNA segment at the insertion site, flanked by the two ME sequences, should contain only the gene of interest and a downstream *attR* site—a hybrid of the *attP* and *attB* motifs. First, we assessed the size of the mini-transposon inserted into the genome in 36 randomly selected antibiotic-sensitive transformants using cPCR (data not shown). In all cases, the DNA fragment matched the expected size after the removal of the Ab^R^ cassette, differing from the fragment sizes obtained from TCJs not transformed with pFNC, used as negative controls. Next, we examined the exact sequence of the *attR* site and nucleotides surrounding the MEs to inspect if Bxb1-int had affected other sequences near the insertion site. We designed oligonucleotides based on the insertion sites found in two *Pp*-CIFR-2 TCJs, targeting the regions of the *thrC* and *phnE* genes affected by the transposase activity. These oligonucleotides, along with an internal primer, were used to amplify the region containing the *attR* sites from the genomes of the corresponding transformants. The amplification products were sent for Sanger sequencing, and both the *attR* site and ME-O matched the expected sequence after the flip-out of the Ab^R^ cassette, underscoring the precision of the Bxb1-int in this process (**Fig. S5**).

Lastly, we evaluated the efficiency of curing pFNC through SacB-mediated counter-selection. This step was performed on the Ab^X^-sensitive (but Ab^Y^-resistant) transformants obtained in the previous phase. Three transformants from each host were inoculated and pooled in LB containing 10% (w/v) sucrose. The cultures were grown overnight to enable the toxic effect of levansucrase to select for cells that no longer retained the plasmid. The cultures were then re-streaked on LB plates to isolate single colonies. Thirty of these single colonies were subsequently re-streaked on LB and LB containing Ab^Y^. All colonies were found to be Ab^Y^-sensitive, indicating that, under these conditions, SacB-mediated curing operated at ∼100% efficiency. An example of the combination *Cn*-CIFR+pFNC-4 is shown in **Fig. S6**. After applying the complete CIFR protocol, the selected clones retained the fluorescent phenotype while being sensitive to both Ab^X^ (used for selecting the initial integration event with pCIFR-X) and Ab^Y^ (adopted for the transformation with pFNC-Y). While the efficiency of the system varied across hosts, with the highest efficiency observed in *P. putida* and *C. necator*, the successful removal of the Ab^R^ cassette, the inability of the integrase to recognize the *attR* scar in the absence of a recombination directionality factor, and the ease of curing the helper plasmid enable the iterative engineering of the same strain using CIFR multiple times in all bacterial hosts.

### 3.6. Expanding the color palette of P. putida in a combinatorial fashion via CIFR

We adopted the CIFR system for integration of expression of different chromoprotein (CP) genes in *P. putida*. CPs are proteins that confer a distinguishable color to corals [87]. We selected the naturally occurring CPs *^ae^*Blue from *Actinia equina*, *^gfas^*Purple from *Galaxea fascicularis*, *^spis^*Pink from *Stylophora pistillata*, and the synthetic *^fw^*Yellow, which is also a fluorescent protein [34, 88, 89]. CPs have not been previously explored as reporters in *P*. *putida*, so we initially expressed the corresponding genes from a plasmid to confirm that these proteins could be produced in this host. The codon-optimized sequences were synthesized and cloned into the expression pSEVA2313 vector under the *P_EM7_* promoter (**Table 2**). Transforming *P. putida* KT2440 with these plasmids led to recognizable phenotypes (**Fig 5a**, left). Next, the four CP genes were cloned into a pCIFR-2 backbone, using the same design as for the expression of the *msfGFP* in the pCIFRs, with the strong promoter *P_14G_* coupled with a BCD2 element to enhance CP production. After transforming *P. putida* with the integration plasmids, strains *Pp*-CIFR-2-*^ae^*Blue, *Pp*-CIFR-2-*^gfas^*Purple, *Pp*-CIFR-2-*^spis^*Pink, and *Pp*-CIFR-2-*^fw^*Yellow were obtained. From each of the four populations, the most intensely colored colony was selected for further analysis. Notably, the expression from a single copy of these genes led to recognizable phenotypes, albeit at a reduced color intensity (**Fig. 5a**, right).

**Fig. 5.**
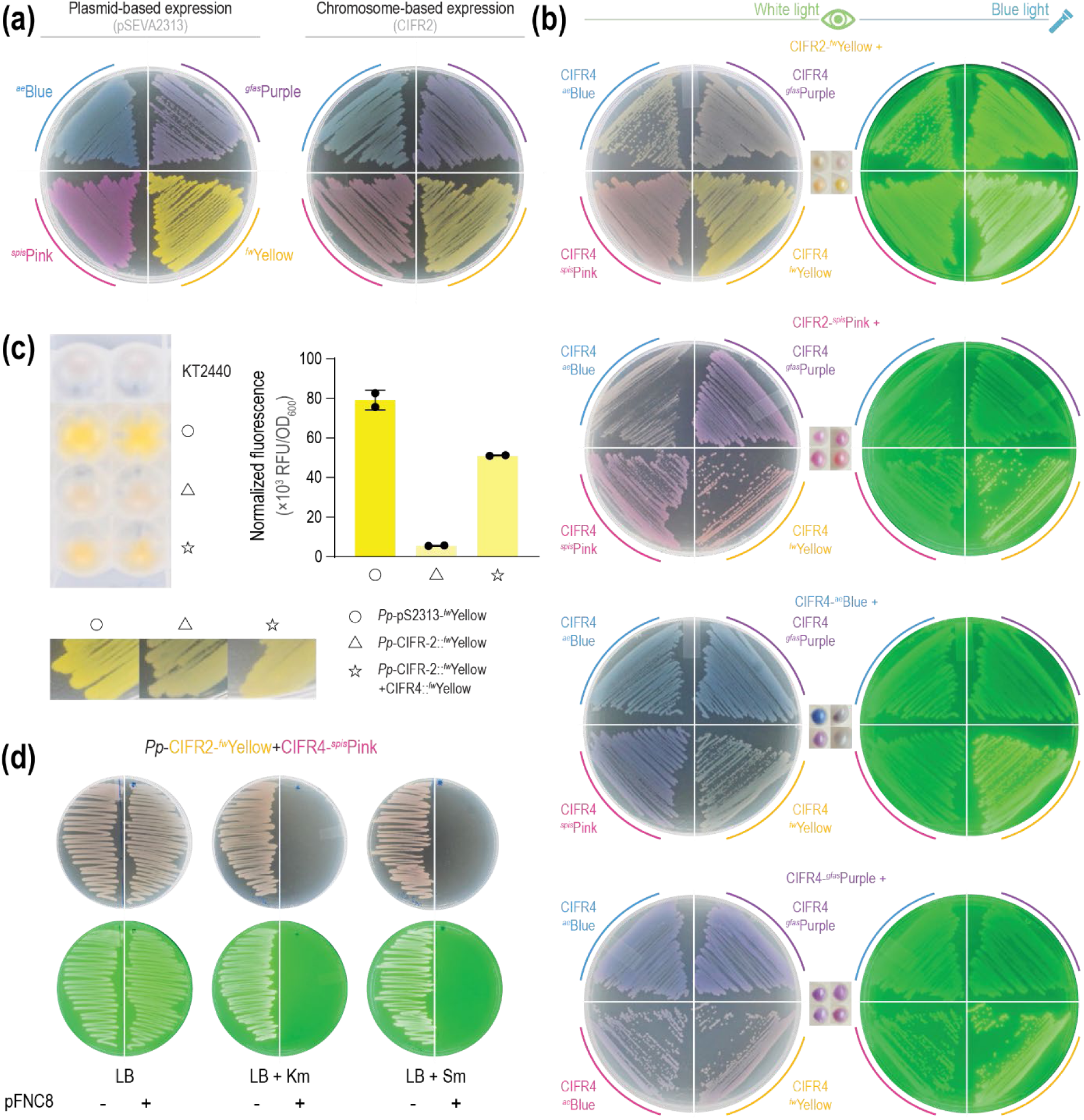
Expression of a library of chromoproteins in *P. putida via* the CIFR system. **(a)** LB plates containing strains of *P. putida* KT2440 carrying the expression cassette for different chromoproteins (i.e. *^ae^*Blue, *^gfas^*Purple, *^spis^*Pink, and *^fw^*Yellow) either on a replicative plasmid (pSEVA2313, left) or integrated in the genome *via* plasmid pCIFR-2 (right). **(b)** Combinatorial screening of double chromoprotein integrations. For each group of plates, the initial integration is indicated at the top-center of the figure; a *P. putida* transconjugant after the initial integration was engineered with a second pCIFR, indicated next to the corresponding streak on the left. The same plates visualized under blue light are shown at the righthand side. In the small square in the center of each set of plates, pellets of the corresponding strains are observed in the same order as in the plates. **(c)** Comparison between the fluorescence intensities of *P. putida* KT2440, expressing *^fw^*Yellow from either a plasmid (○) or a chromosomal integration (△, single; ⋆, double). Cells pelleted in the bottom of a 96-well deep-well plate (top left) are compared to streaks on an LB plate (bottom left). The *^fw^*Yellow fluorescence was also quantified with excitation at 520 nm and emission at 540 nm, and the values were normalized to the optical density at 600 nm (OD_600_, right). Data represent average values from two independent measurements ± standard deviation. **(d)** The strain containing the combinatorial chromoprotein integration performed with plasmids pCIFR-2::*^fw^*Yellow and pCIFR-4:: *^spis^*Pink is resistant to both Km and Sm. After transformation with plasmid pFNC8, both resistances were simultaneously removed.

We adopted the CIFR system for combinatorial engineering of different CP pairs in *P*. *putida*. A second set of plasmids was constructed by cloning the four different CP expression cassettes into vector pCIFR-4 (**Table 2**). The four colored *P. putida* strains previously described were transformed with each of the four pCIFR-4 plasmids. As a result, a total of sixteen transformant combinations were obtained. The phenotypes of these sixteen TCJs displaying pairwise combinations of the CPs exhibited a wide range of color shades and intensities. We observed that combinations starting from *Pp*-CIFR-2-*^spis^*Pink and *Pp*-CIFR-2-*^gfas^*Purple had less color diversity, suggesting that the phenotype conferred by the first CP module was minimally affected by the expression of the second CP gene. Conversely, combinations involving pCIFR-4::*^ae^*Blue resulted in darker shades rather than bluer colors. The combinations using pCIFR-4::*^fw^*Yellow showed the most variations, leading to new colors in all cases. Not surprisingly, the presence of *^fw^*Yellow also conferred fluorescence, in addition to a yellow color, to the strains. The colonies showing the largest difference from the single integrants were selected from each of the sixteen populations and re-streaked (**Fig. 5b**). We named these strains as the combination of the two integrations they carry (e.g., *Pp*-CIFR-2-*^fw^*Yellow+CIFR-4-*^ae^*Blue or *Pp*-CIFR-2-*^fw^*Yellow+CIFR-4-*^gfas^*Purple).

We quantified the effect of the additional copy of the CP gene on the phenotype of strains expressing *^fw^*Yellow, measuring fluorescence (λ_excitation_ = 520 nm/λ_emission_ = 540 nm) of *Pp*-CIFR-2::*^fw^*Yellow and *Pp*-CIFR-2::*^fw^*Yellow+CIFR-4::*^fw^*Yellow. *P. putida* KT2440 transformed with pSEVA2313::*^fw^*Yellow was used as a positive control (**Fig. 5c**), which had the highest maximum normalized fluorescence. This value was 14-fold higher compared to the strain with a single integrated copy of *^fw^*Yellow. The selected strain with two integrated copies of *^fw^*Yellow showed a value of maximum normalized fluorescence 9-fold higher than the one of the single integrant. With two consecutive *^fw^*Yellow integrations, ∼70% of the fluorescence intensity of the plasmid-based expression was achieved. By repeating the CIFR protocol to integrate more copies of the gene, an even higher expression level (therefore, fluorescence intensity) could be achieved, maybe equaling or surpassing the one obtained with replicative plasmid.

The four original strains obtained after the first round of integration and the sixteen strains obtained after the second round of integration were used as templates for arbitrary PCR to determine the insertion site of the different CP-expression cassettes (**Table S8**). Notably, three transposons (in *Pp*-CIFR-2::*^ae^*Blue, *Pp*-CIFR-2::*^gfas^*Purple and *Pp*-CIFR-2::*^ae^*Blue+CIFR-4::*^spis^*Pink) inserted into one of the seven copies of 23S ribosomal RNA present in the genome of *P. putida* KT2440. Ribosomal RNA loci are known to be sites for strong gene expression in *P. putida* [90] and the bias towards highly transcribed spots does not come as a surprise, since the most colorful colonies were picked in this experiment. The distribution of the different genomic location on the chromosome is represented in **Fig. S7**.

*Pp*-CIFR-2::*^fw^*Yellow+CIFR-4::*^spis^*Pink had a different color than the strains with the corresponding single insertions, turning from yellow to pink, and exhibiting fluorescence under blue light excitation. We transformed this strain with pFNC-8 to flip-out the Km and Sm resistance cassettes. One of the antibiotic-sensitive colonies, *Pp*-CIFR::*^fw^*Yellow+CIFR::*^spis^*Pink, retained the color phenotype, while losing the resistance to both antibiotics (**Fig. 5d**). This once again demonstrates the potential of CIFR in building complex genotypes in a rapid and practical fashion.

### 3.7. Carotenoids biosynthesis in P. putida via CIFR-assisted pathway balancing

We then aimed to apply CIFR to address an issue often encountered in metabolic engineering, i.e., pathway balancing. We chose to use the system to introduce a carotenoid biosynthesis pathway in *P. putida* KT2440 that should lead to the production of Zea (**Fig. 6a**). Carotenoids are popular products adopted to test synthetic biology tools, since they are non-peptidic compounds that confers a colored phenotype to producer cells that can be spotted by naked eye and that varies depending on the chemical species produced [35]. Unlike fluorescent proteins and CPs, these compounds require the concerted expression of multiple genes and constitute an ideal proxy to test tools for expression, integration, deletion, and modification of multiple genetic elements. The heterologous production of carotenoids in *P. putida* has been tested by using plasmid expression [91, 92], Tn*7*-assisted site-specific integration [93], and also mini-Tn*5* vectors [94].

**Fig. 6.**
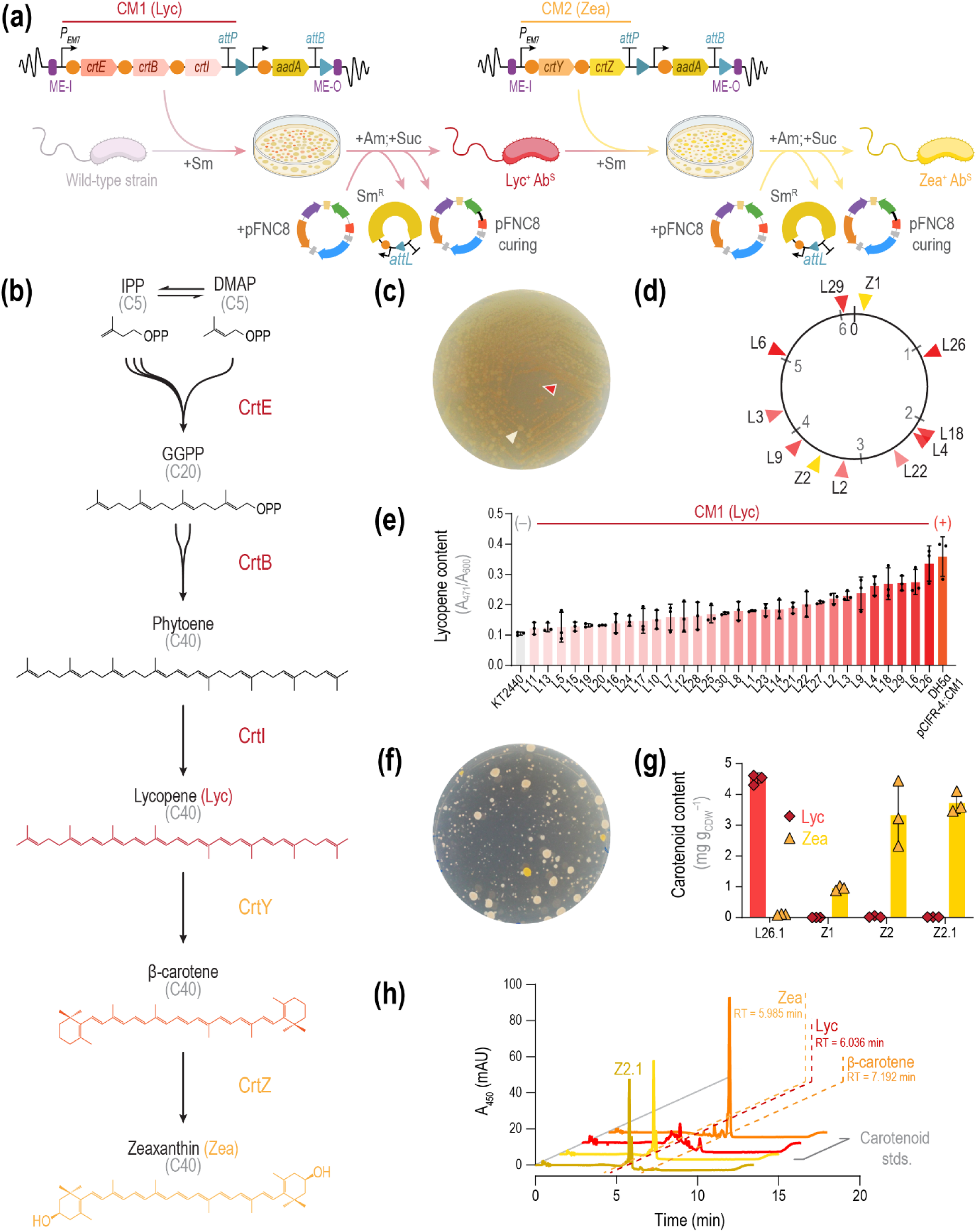
CIFR enables pathway balancing to engineer carotenoid production in *P*. *putida*. **(a)** Modular engineering for establishing zeaxanthin biosynthesis in *P. putida*. The first carotenoid module encompasses the *crtEBI* genes for lycopene (Lyc) biosynthesis from *Pantoea ananatis* (*CM1*), placed under the control of the constitutive *P_EM7_* promoter in plasmid pCIFR-4::*CM1*. This plasmid was used for the first engineering step. After selecting the reddest colony, the Sm resistance was removed by transforming plasmid pFNC-8 in the Lyc^+^ transconjugant. Curing the helper plasmid in a medium with 10% (w/v) sucrose yielded a Lyc-producing, antibiotic-sensitive (Ab^S^) *P. putida* strain. The second carotenoid module, comprising *crtYZ* from *P. ananatis* (*CM2*) to convert Lyc to zeaxanthin (Zea), was integrated using plasmid pCIFR-4::*CM2*. After selecting the yellowest colony, the antibiotic resistance was removed with plasmid pFNC-8 that, after curing, yielded a Zea-producing, Ab^S^ *P. putida* strain. **(b)** Zea biosynthesis pathway from isoprenoid precursors, naturally found in *P. ananatis* and engineered in *P. putida*. The last three compounds are red-(Lyc), orange-(β-carotene), and yellow-colored (Zea). **(c)** *P. putida* streaked on an LB plate containing Sm after transformation with plasmid pCIFR-4::CM1. Both red and white colonies, corresponding to high or low Lyc content, could be observed. **(d)** Genomic integration coordinates of *CM1* in selected red colonies (L) and *CM2* in selected yellow colonies (Z). For the red colonies, the color intensity of the triangle is proportional to the Lyc content. **(e)** Carotenoid content for 30 red colonies isolated from the plate in panel **(c)**, measured as the ratio between A_471_ and OD_600_. *P. putida* KT2440 and *E. coli* DH5α transformed with plasmid pCIFR-4::*CM1* were used as negative and positive controls, respectively. Data represent average values from independent triplicates ± standard deviation. **(f)** L26 colonies upon removal of the resistance cassette, plasmid curing, and transformation with plasmid pCIFR-4::*CM2*. **(g)** Carotenoid content of strain L26.1, either without modifications or engineered with plasmid pCIFR-4::*CM2* integrated in position 1 (Z1) or 2 (Z2). After all engineering steps, the best Zea producer (Z2.1) was isolated as an Ab^S^ and plasmid-free variant of Z2. Carotenoids were quantified by HPLC (detection at λ = 450 nm); data represent average values from three independent measurements ± standard deviation. **(h)** HPLC spectra of selected carotenoids. The spectra of authentic carotenoid standards (Lyc, red; β-carotene, orange; and Zea, yellow; at 50 μg mL^−1^) were recorded at λ = 450 nm, and compared with a sample extracted from Z2.1 (δ). Parts of the figure were adapted from BioRender.com.

The Zea biosynthesis pathway of *Pantoea ananatis* (**Fig. 6b**) starts from the precursors iso pentenyl pyrophosphate (IPP) and dimethylallyl pyrophosphate (DMAPP), produced from pyruvate through the mevalonate pathway [95]. The isoprenoid units are then assembled in the C10 and C15 intermediates geranyl pyrophosphate (GPP) and farnesyl pyrophosphate (FPP), respectively, by farnesyl diphosphate synthase. The geranylgeranyl diphosphate synthase (CrtE) condenses another IPP unit to this last metabolite to yield geranylgeranyl diphosphate (GGPP). Two GGPP units are then condensed together in phytoene by the phytoene synthase (CrtB). Phytoene desaturase (CrtI) performs then multiple desaturations on the hydrocarbon skeleton, to yield the red-pigmented Lyc. The two termini of this carotenoid are then cyclized by Lyc cyclase (CrtY) to obtain the orange-colored β-carotene, dehydroxylated to the yellow compound Zea by the carotene hydroxylase (CrtZ). In this study, we aimed to transfer this pathway in *P. putida* using the CIFR system we developed. *P. putida* is already able to produce IPP and DMAPP *via* the methylerythritol phosphate pathway and can also synthesize GPP and FPP [93]. We therefore had to introduce the genes *crtEBIYZ* from *P. ananatis* in its genome to complete the Zea biosynthetic route. To do this, we split the pathway into two modules: CM1 (*crtEBI*) for the biosynthesis of Lyc, and CM2 (*crtYZ*) for its conversion to Zea (**Fig. 6a-b**).

Lyc is a toxic intermediate, and therefore the expression of the two modules should be carefully balanced in order to avoid the accumulation of this metabolite [91, 96, 97]. Since pathway balancing can be difficult to achieve through rational engineering, requiring multiple regulatory elements, e.g., promoters, RBSs, and terminators [9, 98, 99], we decided to use the random integration capacities offered by CIFR to tackle this issue. As demonstrated in this study, CIFR provides varied expression levels for an integrated expression cassette depending on the landing site, while leaving other regulatory elements unchanged. We therefore cloned CM1 and CM2 into pCIFR-4, setting the genes under the control of the constitutive promoter *P_EM7_* and the standard SEVA RBS [69, 100, 101] and obtaining pCIFR-4::*CM1* and pCIFR-4::*CM2*. We then proceeded to transform *P. putida* KT2440 with plasmid pCIFR-4::*CM1*, obtaining many colonies colored displaying different shades of red (**Fig. 6c**). Big white colonies, akin to wild-type *P. putida* KT2440, were also visible on the plates, probably due to the toxicity of Lyc. Indeed, in some TCJs the *crt* genes were mutated, thus escaping the toxic effect of Lyc. We then aimed to identify the best Lyc producer. We selected the 30 reddest colonies from the plates. L1-30, one of these TCJs, was grown overnight and Lyc was extracted from the biomass with acetone (see *Materials and Methods*). We then measured the absorbance at 471 nm of the organic phase extracts and normalized it by the OD_600_, including wild-type *P. putida* KT2440 and *E*. *coli* DH5α carrying pCIFR-4::*CM1* as a negative and positive control, respectively (**Fig. 6e**). L26 was selected as the best Lyc producer and used for further engineering. The transposon insertion site of 9 Lyc producers was determined by arbitrary PCR (**Fig. 6d** and **Table S9**). In L26, CM1 landed in an intergenic region between *PP_1002* and *PP_1003*. This TCJ was transformed with plasmid pFNC-8 to get rid of the Sm^R^ cassette. Subsequently, we eliminated the helper plasmid to obtain an antibiotic sensitive, Lyc-producing *P. putida* strain, named L26.1. This isolate was transformed with plasmid pCIFR-4::*CM2* and three main types of Sm-resistant colonies could be observed (**Fig. 6f**). Some small reddish colonies were present, a sign that the integration of M2 failed to further convert Lyc to other carotenoid forms. Some other colonies were big and white, hinting that these TCJs blocked the Lyc production pathway to avoid toxicity. Finally, two big colonies exhibited an intense yellow color, suggesting that the corresponding TCJs could produce Zea by expressing CM1 and CM2.

We named these TCJs Z1 and Z2 and mapped the insertion on the genome *via* arbitrary PCR (**Table S10**). We then extracted carotenoids from L26.1, Z1, and Z2 and analyzed the chemical nature of all carotenoids in the samples by HPLC (detection at λ = 450 nm; **Fig. 6g**). While L26.1 produced 4.53 ± 0.16 mg g_CDW_^−1^ of Lyc, Z1, and Z2 accumulated 0.94 ± 0.07 and 3.31 ± 1.06 mg g_CDW_^−1^ of Zea, respectively. Since Z2 was the best Zea producer, the CIFR protocol was completed on this TCJ, yielding an Ab^R^-cassette free, Zea-producing *P. putida*. During these operations, the color and morphology of the colony never changed, demonstrating that Zea production is not as toxic for the cells as Lyc production. The resulting strain, Z2.1, produced 3.76 ± 0.34 mg g_CDW_^−1^ of Zea (corresponding to a titer of 39.7 ± 4.4 mg L^−1^). The Zea content of this strain is one order of magnitude higher than that obtained by Loeschcke, Markert [94], where *crtEBIYZ* were integrated as a single operon in the genome of strain KT2440 using a Tn*5* system. Also, the Zea content of the engineered strain reported here is similar to that of a fully optimized *P. putida* strain where *crtEBIYZ* were expressed from a plasmid together with further manipulations to enhance precursor availability and increased tolerance to carotenoid toxicity [91]. In our work, instead, the heterologous genes are expressed in single copy from the chromosome of wild-type *P. putida* KT2440. Moreover, our final strain produces only Zea and does not accumulate any Lyc and β-carotene, the pathway precursors (**Fig. 6h**). We therefore demonstrate the potential of the CIFR system as an optimal tool for balancing modular pathways, where the expression of the different module must be independently regulated. With our system, the best expression level for each module can be tuned depending on the position on the chromosome, and, if it leads to a particular phenotype, it can be screened individually.

## 4. CONCLUSION AND OUTLOOK

In this study, we presented CIFR, a two-plasmid system designed to integrate synthetic modules into the genome of Gram-negative bacteria and subsequently remove the antibiotic resistance marker used for integration through a streamlined, easy-to-implement protocol. We first demonstrated the functionality of the integration vectors (pCIFR-X), which effectively inserted reporter genes at random loci in the genome of three bacteria with biotechnological potential. Our observations indicate that the Tn*5* system has a slight bias toward genomic regions near the chromosome origin of replication. Next, we demonstrated the helper vectors (pFNC-Y) effectiveness in excising the antibiotic resistance marker positioned between *attP* and *attB*. The excision efficiency varied across microbial hosts, and we recommend verifying this parameter when transferring the system to a new host.

We then showcased two applications of CIFR in *P*. *putida* addressing some of the classical challenges in synthetic biology. First, we provide a palette of reporter genes for *Pseudomonas* species that, unlike fluorescent proteins, can be visualized without specialized equipment—simplifying synthetic biology manipulations with this organism. To this end, we expressed novel CP-encoding genes, both individually and in combination. Additionally, we applied CIFR to address pathway balancing in the biosynthesis of complex products requiring multi-step anabolic routes. The genes for Zea production were distributed across two modules that were integrated sequentially. Adjusting expression levels and selecting TCJs with the highest product content after each step led to an engineered strain with biosynthetic capabilities comparable to those previously reported, where extensive optimization of precursor supply, product tolerance, gene expression, and growth conditions was conducted [91].

While we showcased only two applications of the CIFR system, its potential extends far beyond these examples. The MOI is of course not limited to a reporter gene or a biosynthetic gene cluster. In previous studies, Tn*5* has been employed to integrate regulatory modules, creating a library of mutants with inducible expression of various genes or promoterless reporters to identify regulatory nodes within a genome of interest (**Table 1**). Another promising application for CIFR could involve introducing assimilation modules for diverse substrates—e.g., C1 feedstocks, pollutants, lignocellulosic biomass, or plastic-derived monomers—into bacteria that cannot catabolize these compounds but possess other biotechnologically valuable traits [102]. When engineering these pathways, dividing them into modules and applying individual growth-coupled selection for each module has proven to be an effective strategy for direct selection [103, 104]. We believe that combining growth-coupled selection with Tn5 integration creates an even more powerful strategy, as the semi-random integration in the genome adds an additional layer of expression variability for the synthetic module [105]. A Tn*5* insertion library generated with CIFR can be directly cultured on selective media, allowing transconjugants to compete for nutrients. This process selects for the optimal expression level of the synthetic module required for growth under those conditions.

In this work, we demonstrate the functionality of CIFR in *P*. *putida*, *E*. *coli*, and *C*. *necator*. While these species are prominent biotechnology workhorses, the range of bacteria suitable for sustainable bioproduction is broader. Tn*5* transposons have been used to create mutant libraries across diverse organisms spanning the tree of life, including archaea, bacteria, yeast, plants, and even mammalian and human cells [106–108]. We therefore expect CIFR to work in other Gram-negative hosts, providing a broad-host-range synthetic biology tool to speed up metabolic engineering. In the future, the CIFR collection could be expanded to feature other Ab^R^ cassettes (e.g., Tc^R^ or Cm^R^) to increase the compatibility with emerging microbial hosts. Non-antibiotic resistance cassettes (e.g., those conferring resistance to heavy metals) provide an alternative to select for transformants [18]. We expect that these features will be soon incorporated into the CIFR system for wide adoption by the synthetic biology and metabolic engineering community.

## CREDIT AUTHORSHIP CONTRIBUTION STATEMENT

F.F. and P.I.N. conceived the project. F.F., F.L., and P.I.N. designed the experiments. F.F. and F.L. performed the experiments leading to the results described in this article. C.A.V. and M.E.M. helped with cloning some of the plasmids used in this study. L.B.P. and L.A. helped with the HPLC measurements and analytical determinations. F.F. and F.L. analyzed the data. F.F., F.L., and P.I.N. wrote the manuscript, with contributions from all the other authors.

## DATA AND MATERIAL AVAILABILITY

Data will be made available from the corresponding author upon reasonable request. Plasmids pCIFR and pFNC, carrying different antibiotic resistances, are available in AddGene with the IDs 227663 to 227670.

## ACKNOWLEDGEMENTS

The authors thank the other members of the Systems Environmental Microbiology for fruitful discussions and Verónica Ramos-Viana for help with the HPLC analysis. F.F. is the recipient of a fellowship from the Novo Nordisk Foundation as part of the Copenhagen Bioscience Ph.D. Programme, supported through grants NNF0069783 and NNF20CC0035580. The financial support from The Novo Nordisk Foundation through grants NNF20CC0035580, LiFe (NNF18OC0034818) and TARGET (NNF21OC0067996), the Danish Council for Independent Research (SWEET, DFF-Research Project 8021-00039B), and the European Union’s Horizon 2020 Research and Innovation Programme under grant agreement No. 814418 (SinFonia) to P.I.N. is gratefully acknowledged.

## APPENDIX A · SUPPLEMENTARY DATA

Supplementary data to this article can be found online.

