## Supporting Information for "CIFR (*Clone–Integrate–Flip-out–Repeat*): a toolset for iterative genome and pathway engineering of Gram-negative bacteria"

by

Filippo Federici*, Francesco Luppino*, Clara Aguilar-Vilar, Maria Eleni Mazaraki, Lars Boje Petersen, Linda Ahonen, and Pablo I. Nikel‡

*The Novo Nordisk Foundation Center for Biosustainability, Technical University of Denmark, Kongens Lyngby, Denmark*

* These authors contributed equally and should be considered joint first-authors

**Keywords:** Metabolic engineering; Synthetic biology; transposon; *E. coli*; *Pseudomonas*; *C. necator*; Genome engineering

**Running title:** Transposon tools for bacterial genome engineering

‡ **Correspondence to:**

Pablo I. Nikel

*The Novo Nordisk Foundation Center for Biosustainability*

*Technical University of Denmark, Kongens Lyngby, Denmark*

Preprinted on November 8th 2024.

### SUPPLEMENTARY FIGURES

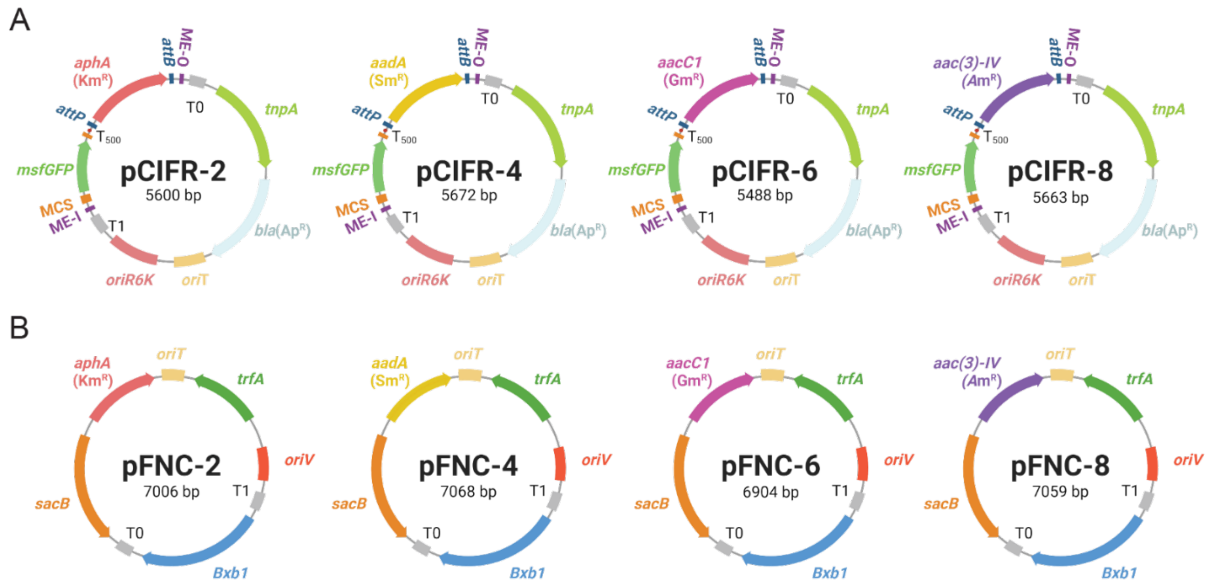


**Fig. S1. Architecture of the pCIFR and pFNC plasmids.** (**a**) pCIFR is a plasmid that contains all the elements to integrate a target sequence in the genome of a Gram-negative host of choice. It is composed by a pir-dependent origin of replication *oriR6K*, an origin of transfer *oriT*, an ampicillin resistance cassette (*bla*), the gene encoding for the transposase *tnpA* and the transposon module, that is transferred to the genome of the host by this same transposase. The transposon module is enclosed by the two identical sequences ME-I and ME-O and contains the gene encoding for msfGFP under the control of strong regulative elements and an antibiotic resistance cassette surrounded by *attP*/*B* sites, that allow its removal after the integration event. pCIFR comes with four different options of removable antibiotic cassette, following the SEVA nomenclature: pCIFR-2: *aphA*, kanamycin resistance; pCIFR-4: *aadA*, streptomycin resistance, pCIFR-6: *aacC1*, gentamicin resistance, pCIFR-8: *aac(3)-IV*, apramycin resistance. Thanks to the presence of unique restriction sites surrounding *msfGFP*, the gene is easily replaceable with a gene or operon of interest using different cloning methods. (**b**) pFNC is a replicative helper plasmid used to remove the resistance marker after performing the integration with pCIFR. pFNC contains the gene encoding for the BxbI phage integrase, performing the recombination between *attP* and *attB*, and the *sacB* gene for easy sucrose-mediated curing of the plasmid. It comes with the same antibiotic resistance cassettes as pCIFR and follows a similar nomenclature. Adapted from BioRender.com.

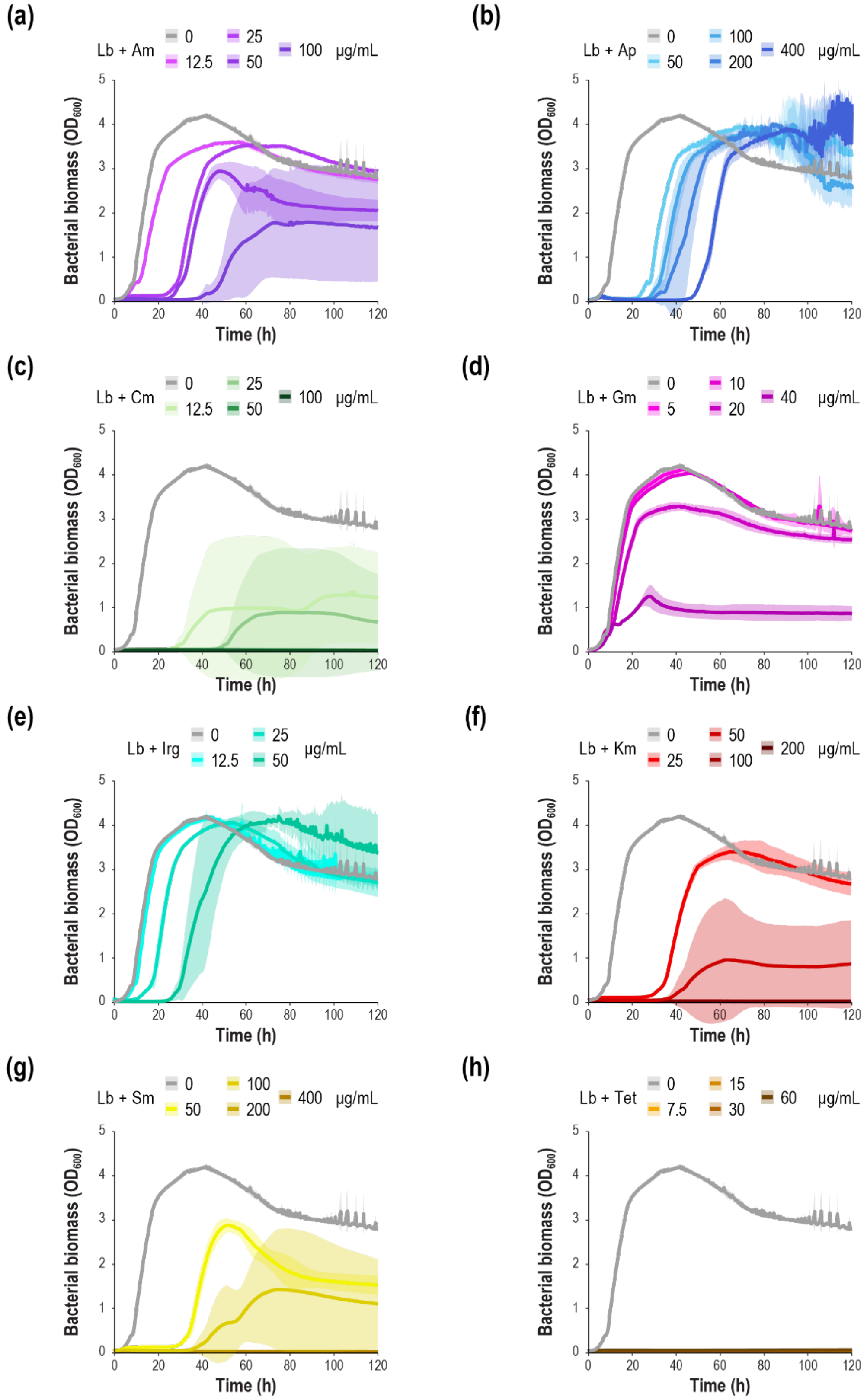


**Fig. S2. Exploration of natural antibiotic tolerance in *C. necator* H16 in liquid LB medium.** *C. necator* H16 was cultivated in a microtiter plate reader in 150 µL of LB medium supplemented with different concentrations of apramycin **(a)**, ampicillin **(b)**, chloramphenicol **(c)**, gentamicin **(d)**, irgasan **(e)**, kanamycin **(f)**, streptomycin **(g)** and tetracycline **(h)**. Optical density at 600 nm ($OD_{600}$) was measured over time. Data represent average values from 2 independent measurements ± standard deviation.

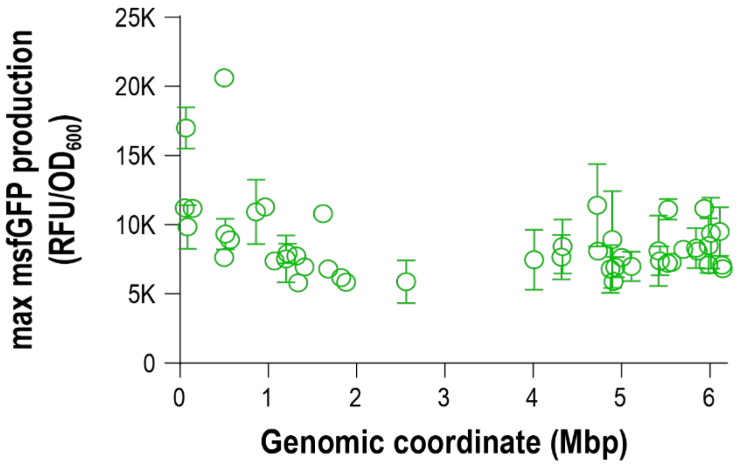


**Fig. S3. Fluorescence intensity and genomic coordinate show no detectable correlation for *msfGFP* integrations in *P. putida* KT2440.** The positional data obtained via arbitrary PCR + sanger sequencing and the fluorescent intensity data of the 47 sequenced clones of *P. putida* KT2440 where *msfGFP* expression cassette was integrated with the CIFR system were plotted against each other. No significative correlation is observed between the two variables. The fluorescence intensity values represent average values from 2 independent measurements ± standard deviation.

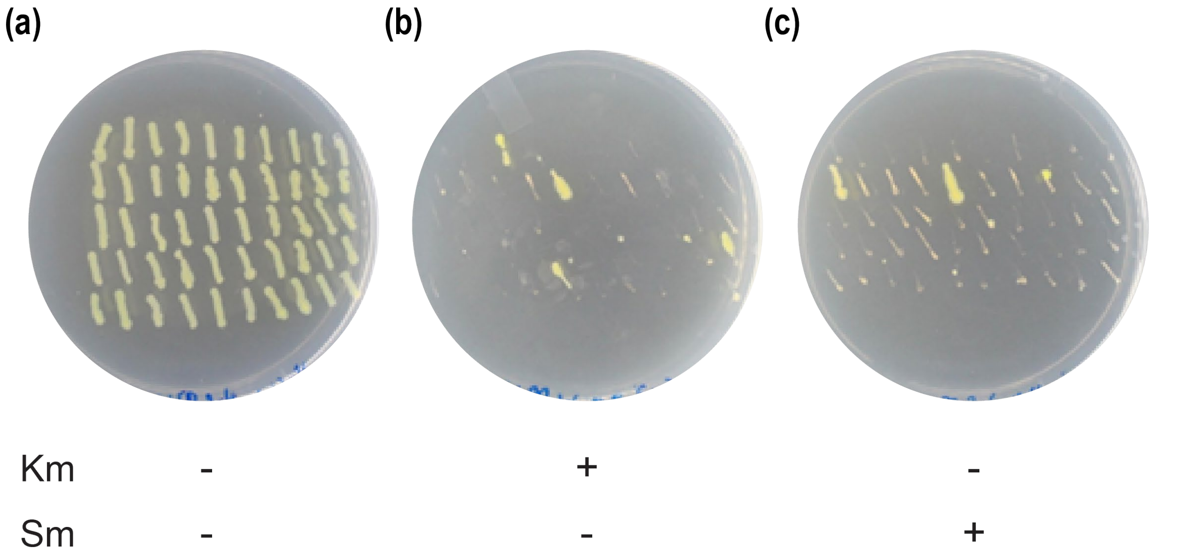


**Fig. S4. pFNC can flip out two antibiotic resistance cassette at the same time.** *P. putida* KT2440 was transformed with pCIFR-2 and pCIFR-4 consecutively. From this double transconjugants, three pools of 5-7 6 colonies were inoculated and transformed with plasmid pFNC-8. Single colonies were isolated, and 50 clones from each transformation were replica-plated on LB plates **(a)**, LB + Km plates **(b)** and LB + Sm plates **(c)**. The efficiency of contextual removal of two antibiotic cassettes mediated by pFNC-Y was determined by dividing the number of colonies sensitive to both antibiotics by the total number of colonies tested. The figure shows an example for one of the three replicates.

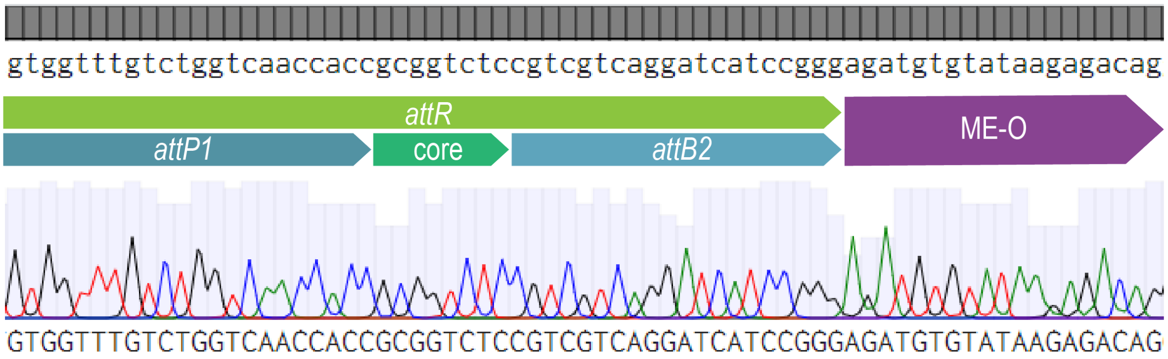


**Fig. S5. Sanger sequencing of the scar left by the CIFR system after removing the kanamycin resistance cassette in *Pp*-CIFR-2 c3.** On the top of the picture, the expected sequence. On the bottom, the chromatograms obtained from the forward and the reverse strand of the amplicon. The *attR* sequence, with segments deriving from *attP* and *attB*, and the 3' mosaic element ME-O are highlighted.

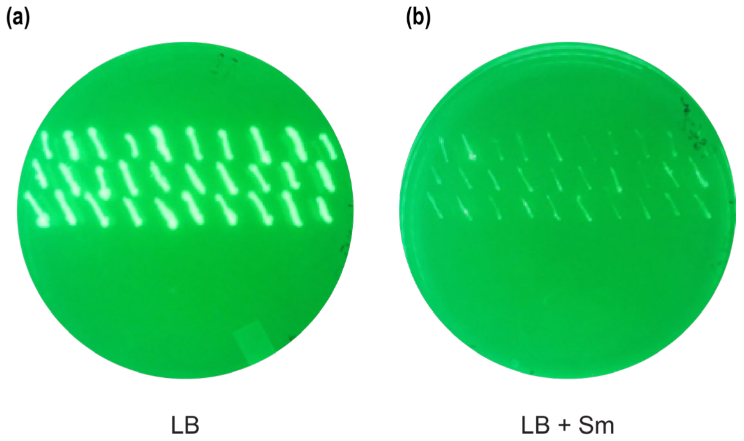


**Fig. S6. Curing efficiency of pFNC-4 in *C. necator* H16.** After transformation of *Cn*-CIFR-2 with pFNC-4, some clones were inoculated in LB + 10% (w/v) sucrose and streaked on an LB plate. The day after, 30 colonies were replica plated on an LB plate (a) and on an LB + Sm plate (b). For *C. necator*, as well as for *P. putida* and *E. coli* (not shown), the SacB-mediated curing efficiency was 100%. The plates are shown under blue light.

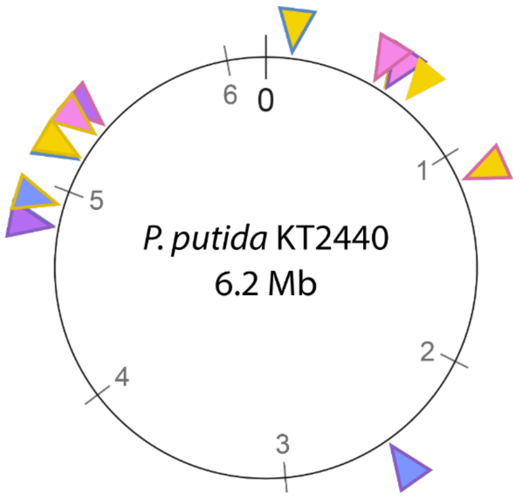


**Fig. S7. Map of the transposon insertion sites of pCIFR harboring chromoprotein expressing cassettes in *P. putida* KT2440.** After selecting the most intensely colored colony for every population of transconjugant transformed with one or two pCIFRs containing a chromoprotein (CP) expression module, an arbitrary PCR was performed on them and sent for Sanger sequencing. Of the four strains expressing only one chromoprotein, only for *Pp*-CIFR-2::$^{fw}$Yellow the insertion site could be specified (yellow triangle with no border), because the other fell into sequences that are repeated through the chromosome. On top of these four strains, a second integration was performed, obtaining 16 additional strains. The position of the second insertion is visible in the map for 13 of these strains, with the filling of the triangle indicating the color of the chromoprotein integrated with this last transformation, and the border indicating the previous color of the strain on which this was performed.

### SUPPLEMENTARY METHODS

*Details on the cloning of the plasmids used in this study*

The cloning of pCIFR-2/4/6 was performed in two steps. First, the *attP* and *attB* cassettes were placed surrounding the antibiotic resistance cassette in pBAMD1-2/4/6 using primers Tn5R_BB_U_fw, Tn5R_BB_U_rv, Tn5R_AbR_U_fw and Tn5R_AbR_U_rv, obtaining pBAMD1-2/4/6R. These plasmids were then linearized using primers pBAMD_operon_U_fw and pBAMD_operon_U_rv and $P_{14g}$(BCD2)→*msfGFP* was amplified using p14g_U_fw and msfGFP_U_rv from pSNW2 and inserted into the MCS (Volke et al., 2020). To obtain pCIFR-8, the Km$^R$ cassette of pCIFR-2 was exchanged with the Am$^R$ cassette from pSEVA88c1 using Am_CIFR_bb_U_fw and Am_CIFR_bb_U_rv (for the backbone) and Am_CIFR_U_fw and Am_CIFR_U_rv (for the cassette)(García-Gutiérrez et al., 2020).

pFNC-6 was also cloned in two steps. First, the *sacB* counterselection cassette from pMBEC2 was amplified with SacB_gadget_U_fw and SacB_gadget_U_rv and placed in the gadget position of pSEVA6213s (linearized with pSEVA_Gadget_ins_U_fw and Chrmp_U_rv), obtaining pSEVA6213s-sacB (Silva-Rocha et al., 2013; Wirth et al., 2020). Then, the BxbI phage integrase gene, amplified with BxBI_rbs_U_fw and BxbI_U_rv from pBad-Int Set Flipper, was cloned in the MCS of pSEVA6213s-sacB, linearized with pS62xSac_BB_U_fw and pEM7_U_rv, to obtain pFNC-6 (Bonnet et al., 2012). pFNC-2/4/8 are derivatives of pFNC-6 were the Gm$^R$ cassette has been exchanged with Km$^R$, Sm$^R$ or Am$^R$ (using primers SEVA_AbEX_BB_U_fw + SacB_gadget_U_rv, SEVA_AbEX_Km_U_fw + SEVA_AbEX_Km_U_rv, SEVA_AbEX_Sm_U_fw + SEVA_AbEX_Sm_U_rv and SEVA_AbEX_Am_U_fw + SEVA_AbEX_Am_U_rv, to amplify the backbone and the three cassettes Km$^R$, Sm$^R$ and Am$^R$, respectively).

To clone the CP-encoding genes into pSEVA2313 they were amplified from the synthesized DNA with oligos Chrmp_U_fw and Chrmp_U_rv (Wirth et al., 2020). The genes were synthesized already equipped with RBS and terminator in order to use the same set of primers for all of them. The backbone of pSEVA2313 was linearized with Chrmp_BB_U_fw and Chrmp_BB_U_rv. The CP expression units (promoter, RBS, CDS) were then transferred to pCIFR-2/4 using the primers p14G_U_fw + Chrmp_int_U_rv and Chrmp_CIFR_BB_U_fw + Chrmp_CIFR_BB_U_rv for the expression unit and the backbone, respectively.

To clone pCIFR-4::CM1, the genes *crtEBI*, with the respective ribosome binding sites (RBSs), were amplified using CrtEBI_U_fw and CrtEBI_U_rv from pPS1·CRT and inserted instead of $P_{14g}$(BCD2)→*msfGFP* in pCIFR-4, linearized with Crt_CIFR_BB_U_fw and Crt_CIFR_BB_U_rv (Sánchez-Pascuala et al., 2019). In the cloning, the $P_{EM7}$ promoter was amplified from pSEVA6231s with primers PEM7_U_fw and PEM7_U_rv2 and placed in front of the operon (Wirth et al., 2020). pCIFR-4::CM2 was cloned in a similar manner, using the same fragments for backbone and promoter and adding *crtY* (amplified with CrtY _U_fw and CrtYZ_rbs_U_rv from pPS1·CRT) and *crtZ* (amplified from the ordered GeneBlock using primers CrtYZ_rbs_U_fw and CrtZ_U_rv).

#### SUPPLEMENTARY TABLES

**Table S1.** List of oligonucleotides used in this study.

| Name | DNA sequence (5'→3') | Use |
|---|---|---|
| **Tn5R_BB_U_fw** | ACGGCGGUCTCCGTCGTCAGGATCATCCGGGAGATGTGTATAAGAGACAGACTAG | Inserting *attP*/*attB* sites in pBAMD1-2/4/6 |
| **Tn5R_BB_U_rv** | ACCGCGGUGGTTGACCAGACAAACCACACAGAAAAGCCCGCCTTT | |
| **Tn5R_AbR_U_fw** | ACCGCGGUCTCAGTGGTGTACGGTACAAACCCAATTTAAATTTGTGTCTCAAA | |
| **Tn5R_AbR_U_rv** | ACCGCCGUCGTCGACAAGCCGGCCTCAGAGATTTTATGCTTGTAAACC | |
| **pBAMD_operon_U_fw** | ATAATCUAGAGTCGACCTGCAGGCATGC | Inserting $P_{14g}$(BCD2)→*msfGFP* in the MCS of pBAMD1-2/4/6R |
| **pBAMD_operon_U_rv** | ATGGGCGGAUCCCCGGGTACCGAGCTCGAA | |
| **p14g_U_fw** | ATCCGCCCAUTGACAAGGCTCTCGCGGC | |
| **msfGFP_U_rv** | AGATTAUTTGTAGAGTTCATCCATG | |
| **Am_ClFR_U_fw** | ATTTAAAUCTCACGGTAACTGATGCCG | Exchanging $Km^R$ for $Am^R$ in pClFR-2, to obtain pClFR-8 |
| **Am_ClFR_U_rv** | ACAACGGUCTCAGCCAATCGACTGGC | |
| **Am_ClFR_bb_U_fw** | ACCGTTGUCTAATTAATTGCGGACCC | |
| **Am_ClFR_bb_U_rv** | ATTTAAAUTGGGTTTGTACCGTACAC | |
| **SacB_gadget_U_fw** | ATCCAGGGGUCCCCCGGCATCAGAGCAGATT | Inserting *sacB* in the gadget site of pSEVA6213s |
| **SacB_gadget_U_rv** | ATTTAAAUGCAAGACCTAAAATGTGTAAAGG | |
| **pSEVA_Gadget_ins_U_fw** | ATTTAAAUTTGACATAAGCCTGTTCG | |
| **Chrmp_U_rv** | ACCCCTGGAUTCTCACCAATAAAAAAACG | |
| **BxBI_rbs_U_fw** | AACCAAUGGAGTCATGACCATGCCTAGGT | Exchanging *I-sceI* for the BxbI phage integrase gene in the MCS of pSEVA6213s |
| **BxBI_U_rv** | AGCGTTATUACGACATCCCGGTGTGTA | |
| **pS62xSac_BB_fw** | AATAACGCUGATAGTGCTAGTGTAGATCGCTAC | |
| **pEM7_U_rv** | ATTGGTUTAGTTCCTCACCTTGTCGTATTAT | |
| **SEVA_AbEX_BB_U_fw** | ACCGCGGUCCGCGCGTTGTCCTTTTC | Exchanging antibiotic resistances in pFNC-6 to obtain pFNC-2/4/8 |
| **SEVA_AbEX_Km_U_fw** | ATTTAAAUTTGTGTCTCAAAATCTCTGATGTT | |
| **SEVA_AbEX_Km_U_rv** | ACCGCGGUCCAATTAATTATTAGAAAAATTC | |
| **SEVA_AbEX_Sm_U_fw** | ATTTAAAUGAACCTTGACCGAACGCA | |
| **SEVA_AbEX_Sm_U_rv** | ACCGTTGUCTTATTTGCCGACTACCT | |
| **SEVA_AbEX_Am_U_fw** | ATTTAAAUCTCACGGTAACTGATGCCGTAT | |
| **SEVA_AbEX_Am_U_rv** | ACCGCGGUCTCAGCCAATCGACTGG | |
| **Chrmp_U_fw** | ACAAAAGCUTAGGAGGAAAAACATATG | Cloning CP-encoding genes into the MCS of pSEVA2313 |
| **Chrmp_BB_U_fw** | ATCCAGGGGUCCCCAATAATTACGATTTAA | |
| **Chrmp_BB_U_rv** | AGCTTTTGUCGTATTATACTATGCCGA | |
| **Chrmp_int_U_rv** | AGACTGGAUTCTCACCAATAAAAAAACG | Cloning CP-encoding genes into the MCS of pClFR2/4 |
| **Chrmp_ClFR_BB_U_fw** | ATCCAGTCUAGAGTCGACCTGCAGGCATGCA | |
| **Chrmp_ClFR_BB_U_rv** | ATTAGAAAACCUCCTTAGCATGATTAAGAT | |

| | | |
|---|---|---|
| **Crt_CIFR_BB_U_fw** | ATCTAGAGUCGACCTGCAGGCATGCAAGCT | *Cloning crtEBI and crtYZ of P. anantis into pCIFR-4* |
| **Crt_CIFR_BB_U_rv** | ATTGTCACGGAUCCCCGGGTACCGAGCTCG | |
| **pEM7_U_fw** | ATCCGTGACAAUTAATCATCGGCATAGTATATCGG | |
| **pEM7_U_rv2** | ATATGTUTTTCCTCCTCCTAGGGGTTT | |
| **CrtEBI_U_fw** | AACATAUGACGGTCTGCGCAAAAAAAC | Cloning *crtEBI* of *P. anantis* into pCIFR-4 |
| **CrtEBI _U_rv** | CTCTAGAUTATATCAGATCCTCCAGCATCAA | |
| **CrtY_U_fw** | AACATAUGCAACCGCATTATGATCTGATTCT | Cloning *crtYZ* of *P. anantis* into pCIFR-4 |
| **CrtYZ_rbs_U_rv** | ATGTTTTUCCTCCTCCTAGGTTAACGATGAGTCGTCATAA | |
| **CrtYZ_rbs_U_fw** | AAAAACAUATGTTGTGGATCTGGAACGC | |
| **CrtZ_U_rv** | ACTCTAGAUCATTTTCCGCTAGCTGGCTCAT | |
| **attB-Ext-F** | GGCCGGCTTGTCGACGACGGCGGTCTC | First round of arbitrary PCR |
| **ARB6** | GGCACGCGTCGACTAGTACNNNNNNNNNNACGCC | |
| **Sm_Ext_F** | GGCGAGATCACCAAGGTAGTCGGCAAATAA | First round of arbitrary PCR, specific for pCIFR-4 |
| **attB-Int-F** | GCGGTCTCCGTCGTCAGGATCATCCGG | Second round of arbitrary PCR |
| **ARB2** | GGCACGCGTCGACTAGTAC | |

**Table S2.** List of synthetic gene blocks used in this study. All the sequences were codon optimized for P. putida.

| Name | DNA sequence (5'→3') |
|---|---|
| **RBS/$^{ae}$Blue/T0** | AAGCTTAGGAGGAAAAACATATGGCTAGCCTGGTTAAGAAAGACATGTGCATCAAGATGACG<br>ATGGAAGGAACAGTAAACGGCCACCACTTCAAGTGCGTCGGCGAGGGCGAAGGCAAGCCGTT<br>CGAAGGCACCCAGGTGGAGAAGATCCGCATCACTGAAGGTGGGCCCCTCCCATTCGCGTACG<br>ACATCCTGGCCCCATGCTGCATGTATGGCAGCAAAACCTTCATTAAGCACGTGTCGGGTATC<br>CCGGACTACTTTAAGGAGTCGTTCCCTGAGGGCTTTACCTGGGAACGCACCCAGATCTTCGA<br>GGATGGCGGCTATCTCACCATCCACCAGGACACGAGCTTGCAGGGTAACAATTTCATTTTCA<br>AGGTGAACGTCATCGGTGCCAACTTCCCTGCCAACGGCCCCGTGATGCAGAAAAAGACGGCA<br>GGCTGGGAACCGTGCGTCGAGATGCTTTACCCGCGGGACGGCGTCCTGTGTGGCCAGAGCCT<br>GATGGCCCTGAAGTGCACCGATGGCAACCACCTGACCTCCCACCTGCGCACCACCTACCGTT<br>CCCGCAAGCCCTCGAATGCAGTGAACATGCCGGAATTTCATTTCGGGGACCATCGCATAGAG<br>ATCTTGAAAGCCGAACAAGGTAAGTTCTATGAACAATACGAGTCAGCGGTGGCCCGTTACTG<br>TGAGGCGGCCCCGAGTAAACTGGGGCATCACTAATGACTAGTCTTGGACTCCTGTTGATAGA<br>TCCAGTAATGACCTCAGAACTCCATCTGGATTTGTTCAGAACGCTCGGTTGCCGCCGGGCGT<br>TTTTTATTGGTGAGAATCCAG |
| **RBS/$^{fw}$Yellow/T0** | AAGCTTAGGAGGAAAAACATATGACCGCACTGACTGAAGGCGCCAAGCTCTTCGAGAAAGAA<br>ATCCCATACATCACTGAGCTTGAGGGGGACGTGGAGGGTATGAAGTTTATCATCAAGGGCGA<br>GGGTACAGGGGACGCGAGCGTCGGAAAGGTGGATGCTCAGTTCATTTGTACCACGGGCGACG<br>TTCCGGTCCCGTGGAGCACCTTGGTCACCACGCTGACTTATGGGGCTCAGTGCTTCGCCAAG<br>TACCCGCGCCACATAGCGGACTTCTTCAAAAGCTGTATGCCGGAGGGGTACGTCCAAGAGCG<br>CACCATCACCTTCGAGGGTGACGGCGTGTTCAAGACCCGTGCGGAAGTCACCTTTGAAAACG<br>GCAGCGTGTACAACCGGGTAAAGCTGAACGGCCAGGGTTTCAAGAAGGACGGCCACGTGCTG<br>GGCAAGAACCTGGAGTTCAACTTCACCCCTCACTGCTTGTACATTTGGGGTGACCAGGCGAA<br>CCATGGCCTGAAGAGCGCGTTCAAGATCATGCATGAGATCACCGGCTCCAAAGAGGATTTCA<br>TCGTTGCCGATCACACCCAAATGAACACCCCCATCGGCGGGGGGCCCGTGCACGTGCCAGAG<br>TACCACCACATCACGTATCATGTTACCTTGAGCAAGGACGTCACCGACCACCGGGACCATTT<br>GAACATCGTGGAGGTGATCAAGGCCGTAGACCTGGAGACGTACCGGTGATAAACTAGTCTTG<br>GACTCCTGTTGATAGATCCAGTAATGACCTCAGAACTCCATCTGGATTTGTTCAGAACGCTC<br>GGTTGCCGCCGGGCGTTTTTTATTGGTGAGAATCCAG |
| **RBS/$^{gfas}$Purple/T0** | AAGCTTAGGAGGAAAAACATATGTCGGTGATCGCCAAGCAGATGACCTACAAAGTCTATATG<br>TCGGGTACTGTGAACGGCCATTATTTTGAAGTGGAGGGTGACGGGAAGGGCAAGCCGTATGA<br>AGGCGAACAGACCGTTAAACTTACCGTCACGAAGGGCGGCCCATTGCCGTTCGCATGGGATA<br>TTCTGAGCCCCCAGTCCCAATATGGCAGCATCCCATTCACGAAGTACCCGGAGGACATCCCG<br>GACTACGTGAAGCAGAGTTTTCCGGAGGGTTACACCTGGGAACGAATCATGAACTTCGAAGA<br>CGGCGCCGTCTGCACCGTGAGCAACGACAGCTCTATCCAGGGCAATTGCTTCATCTACCATG<br>TCAAGTTCTCAGGTCTGAACTTCCCGCCGAACGGCCCGGTGATGCAGAAGAAGACCCAGGGC<br>TGGGAGCCTAATACAGAACGGCTGTTCGCCCGCGATGGAATGTTGATCGGCAACAATTTCAT<br>GGCTTTGAAGCTGGAGGGCGGTGGCCACTATCTGTGCGAGTTTAAGAGCACCTACAAGGCGA<br>AAAAGCCCGTTAAGATGCCCGGCTATCATTACGTGGATCGTAAGCTGGACGTTACCAACCAC<br>AACAAGGACTACACCTCCGTCGAACAGTGCGAGATTTCGATCGCCCGCAAGTCGGTGGTTGC<br>CTAATAG/ACTAGTCTTGGACTCCTGTTGATAGATCCAGTAATGACCTCAGAACTCCATCTG<br>GATTTGTTCAGAACGCTCGGTTGCCGCCGGGCGTTTTTTATTGGTGAGAATCCAG |
| **RBS/$^{spis}$Pink/T0** | AAGCTTAGGAGGAAAAACATATGTCGCACTCCAAGCAGGCCCTGGCAGACACAATGAAAATG<br>ACCTGGCTGATGGAAGGGTCGGTCAACGGGCACGCCTTTACCATCGAAGGCGAAGGTACAGG<br>CAAACCGTACGAAGGCAAGCAGAGTGGCACCTTCCGTGTGACCAAAGGCGGTCCCCTGCCGT<br>TTGCCTTTGACATCGTCGCTCCTACCCTGAAGTACGGCTTCAAATGCTTCATGAAGTACCCC<br>GCGGATATACCCGACTACTTTAAGCTGGCCTTCCCCGAAGGCCTCACGTACGACCGCAAAAT<br>TGCGTTTGAGGACGGCGGATGCGCCACCGCCACGGTCGAAATGAGCCTAAAGGGCAACACCC<br>TCGTGCATAAAACGAACTTCCAGGGCGGAAATTTCCCGATTGACGGGCCGGTGATGCAGAAG<br>CGCACCCTGGGCTGGGAGCCGACCTCTGAGAAGATGACTCCGTGCGACGGTATCATCAAAGG<br>CGACACCATCATGTACCTGATGGTAGAAGGCGGCAAGACTCTGAAATGTAGGTACGAAAACA<br>ATTACCGCGCCAACAAGCCAGTGCTGATGCCGCCTAGCCACTTCGTGGATCTCCGCCTCACC<br>CGCACAAACCTGGACAAAGAAGGTCTGGCGTTCAAACTGGAGGAATATGCCGTCGCCCGCGT<br>GCTGGAAGTGTGATGA/ACTAGTCTTGGACTCCTGTTGATAGATCCAGTAATGACCTCAGAA |

| | |
|---|---|
| | CTCCATCTGGATTTGTTCAGAACGCTCGGTTGCCGCCGGGCGTTTTTTATTGGTGAGAATCC AG |
| ***crtZ*** | ATGTTGTGGATCTGGAACGCCCTGATCGTTTTCGTCACCGTGATCGGCATGGAAGTGATTGC CGCCCTGGCCCACAAGTACATCATGCACGGCTGGGGTTGGGGCTGGCACCTGTCCCATCATG AACCGCGTAAGGGTGCGTTCGAAGTCAACGACCTGTACGCCGTGGTATTTGCTGCACTCTCG ATCCTGCTGATCTATCTGGGCAGTACCGGCATGTGGCCGCTGCAGTGGATCGGCGCCGGTAT GACCGCGTACGGACTGCTGTATTTCATGGTGCACGACGGGCTGGTGCACCAACGCTGGCCTT TCCGCTACATTCCACGCAAGGGCTACTTGAAACGGTTGTACATGGCCCACCGTATGCATCAC GCCGTCCGCGGCAAGGAGGGTTGCGTGTCGTTCGGCTTCCTCTATGCGCCGCCCCTGAGCAA ACTTCAGGCGACGCTCCGGGAGCGCCATGGCGCTCGCGCCGGAGCAGCCCGTGACGCGCAAG GTGGCGAAGATGAGCCAGCTAGCGGAAAATGA |

**Table S3.** Exploration of natural antibiotic tolerance in *C. necator* H16 in liquid and solid LB medium.

| **Antibiotic** | **1x concentration [µg/mL]** | **Growth in liquid LB medium at antibiotic concentration =** | | | | **Growth in LB + 1,5% (w/v) agar plate at conc. 1x** |
|---|---|---|---|---|---|---|
| | | **4x** | **2x** | **1x** | **½x** | |
| Am | 25 | ++ | +++ | +++ | +++ | ++ |
| Ap | 100 | +++ | +++ | +++ | +++ | ++ |
| Cm | 25 | - | - | + | + | - |
| Gm | 10 | ++ | +++ | +++ | +++ | +++ |
| Irg | 25 | / | +++ | +++ | +++ | / |
| Km | 50 | - | - | + | +++ | + |
| Sm | 100 | - | - | + | ++ | - |
| Tet | 15 | - | - | - | - | / |

*C. necator* H16 was cultivated either in 150 µL of LB medium in a microtiter plate reader or on LB + 1,5% agar plates. The media was supplemented with different concentrations of apramycin (Am), ampicillin (Ap), chloramphenicol (Cm), gentamicin (Gm), irgasan (Irg), kanamycin (Km), streptomycin (Sm) and tetracyclin (Tet). The concentrations tested in liquid media were always multiples (1/2x, 1x, 2x, 4x) of the concentration tested in solid medium, and reported in the second column from the left. In the microtiter plate reader, optical density at 600 nm ($OD_{600}$) was measured over time. On plates, growth was qualitatively evaluated. Legend: +++: final $OD_{600}$ similar to the one in LB without antibiotics/ same colonies on plate as on a plate without antibiotics; ++: final $OD_{600}$ < 50% of that in LB without antibiotic/ less colonies on plate than on a plate without antibiotics; + only some replicates could grow and with final OD < 50% of that in LB without antibiotic / very few colonies on plate; -: no replicates could grow in liquid LB/ no colonies on plate; /: not tested. Qualitative observation made from 2 independent replicates.

**Table S5.** Transposon insertion sites of pCIFR in *P. putida* KT2440.

| Colony | Genome coordinate (bp) | PP number | Strand | Gene name, putative function | max msfGFP production (RFU/$OD_{600}$) |
|---|---|---|---|---|---|
| *Pp*-CIFR-2 #1 | 6,146,885 | 5392 | - | WD40/YVTN repeat-containing protein | 6,823 ± 5 |
| *Pp*-CIFR-2 #2 | 5,985,810 | 5245 | - | AraC family transcriptional regulator | 7,087 ± 4 |
| *Pp*-CIFR-2 #3 | 1,676,311 | 1471 | + | *thrC*, threonine synthase | 6,800 ± 7 |
| *Pp*-CIFR-2 #4 | 964,366 | 0826 | + | *phnE*, phosphonate ABC transporter permease | 11,271 ± 32 |
| *Pp*-CIFR-2 #6 | 5,433,630 | 4772 | - | *hrpB*, ATP-dependent helicase HrpB | 7,383 ± 1,051 |
| *Pp*-CIFR-2 #7 | 4,723,403 | 4179 | + | *htpG*, chaperone protein | 11,407 ± 2,983 |
| *Pp*-CIFR-2 #8 | 4,732,783 | 4188-4189 | - | intergenic region | 8,092 ± 86 |
| *Pp*-CIFR-2 #9 | 5,523,618 | 4857 | - | *yhjG*, uncharacterized protein | 7,212 ± 81 |
| *Pp*-CIFR-2 #10 | 6,139,985 | 5386 | + | *cusB*, RND family copper transporter membrane fusion protein | 7,079 ± 28 |
| *Pp*-CIFR-2 #11 | 500,135 | 0411 | + | polyamine ABC transporter ATP-binding protein | 7,632 ± 23 |
| *Pp*-CIFR-2 #12 | 5,698,362 | 5001 | + | *hslU*, protease HslVU ATPase subunit | 8,220 ± 102 |
| *Pp*-CIFR-2 #15 | 5,987,144 | 5246 | - | *kefB-III*, glutathione-regulated potassium/H+ antiporter | 8,479 ± 1,992 |
| *Pp*-CIFR-4 #1 | 5,861,266 | 5136 | + | ABC transporter permease | 8,096 ± 120 |
| *Pp*-CIFR-4 #2 | 500,129 | 0411 | + | polyamine ABC transporter ATP-binding protein | 20,618 ± 466 |
| *Pp*-CIFR-4 #3 | 1,315,926 | 1147-1148 | + | intergenic region | 7,732 ± 442 |
| *Pp*-CIFR-4 #4 | 1,202,006 | 1052 | + | *xcpW*, type II secretion pathway protein | 7,532 ± 1,681 |
| *Pp*-CIFR-4 #5 | 1,338,757 | 1163 | - | FAD-binding oxidoreductase | 5,807 ± 296 |
| *Pp*-CIFR-4 #8 | 1,070,923 | 0928 | - | K+-dependent Na+/Ca+ exchanger-like protein | 7,378 ± 106 |
| *Pp*-CIFR-4 #9 | 5,569,797 | 4899 | - | *nnrD,* ADP-dependent (S)-NAD(P)H-hydrate dehydratase | 7,275 ± 425 |
| *Pp*-CIFR-4 #10 | 4,331,479 | 3803 | + | *pfvE*, ABC-type Fe2+ transporter permease component | 8,421 ± 1,939 |
| *Pp*-CIFR-4 #11 | 6,010,514 | 5263 | - | GGDEF domain-containing protein | 9,381 ± 2,571 |

| | | | | | |
|---|---|---|---|---|---|
| *Pp*-CIFR-4 #12 | 5,417,458 | 4757 | + | *gudD*, bifunctional D-glucarate dehydratase/L-idarate epimerase | 8,116 ± 2,533 |
| *Pp*-CIFR-4 #15 | 4,910,739 | 4320 | + | cycH, cytochrome c-type biogenesis protein | 5,912 ± 125 |
| *Pp*-CIFR-6 #1 | 5,843,583 | 5122 | - | GMC family oxidoreductase | 8,299 ± 1,446 |
| *Pp*-CIFR-6 #3 | 329,772 | 0271 | - | *gltR-I*, two-component system response regulator | 11,604 ± 129 |
| *Pp*-CIFR-6 #4 | 66,420 | 0056 | + | *betA-I*, choline dehydrogenase | 16,979 ± 1,502 |
| *Pp*-CIFR-6 #5 | 4,320,053 | 3790-3791 | + | intergenic region | 7,659 ± 1,603 |
| *Pp*-CIFR-6 #7 | 4,928,946 | 4338 | + | *cheA*, chemotaxis histidine kinase CheA | 6,917 ± 746 |
| *Pp*-CIFR-6 #8 | 84,195 | 0073-0074 | + | Intergenic region | 9,854 ± 1,596 |
| *Pp*-CIFR-6 #9 | 5,110,485 | 4495 | + | *aroP-II*, aromatic amino acid transport protein | 6,989 ± 1,063 |
| *Pp*-CIFR-6 #10 | 566,326 | 0482 | + | *bfr-I*, bacterioferritin 1 | 8,910 ± 416 |
| *Pp*-CIFR-6 #11 | 1,408,385 | 1230 | + | hypothetical protein | 6,952 ± 393 |
| *Pp*-CIFR-6 #12 | 4,008,030 | 3535-3536 | - | gamma-glutamyltransferase | 7,470 ± 2,161 |
| *Pp*-CIFR-6 #13 | 2,558,754 | 2242 | - | *fepA*, ferric enterobactin transport system outer membrane subunit | 5,876 ± 1,549 |
| *Pp*-CIFR-6 #14 | 4,895,723 | 4304 | + | VIC family cation transporter | 8,937 ± 3,509 |
| *Pp*-CIFR-6 #15 | 143,002 | 0134-5428 | - | intergenic region | 11,174 ± 536 |
| *Pp*-CIFR-6 #16 | 52,731 | 0046 | - | *opdT-I,* tyrosine-specific outer membrane porin D | 11,222 ± 19 |
| *Pp*-CIFR-8 #4 | 1,218,953 | 1064 | + | alpha/beta family hydrolase | 7,922 ± 714 |
| *Pp*-CIFR-8 #5 | 858,928 | 0740 | - | MerR family transcriptional regulator | 10,929 ± 2,340 |
| *Pp*-CIFR-8 #6 | 512,875 | 0424 | - | *crp*, DNA-binding transcriptional dual regulator | 9,332 ± 1,1120 |
| *Pp*-CIFR-8 #7 | 6,113,159 | 5364 | - | *clsA*, cardiolipin synthase | 9,514 ± 1,746 |
| *Pp*-CIFR-8 #9 | 1,881,700 | 1689 | - | *fadL*, long-chain fatty acid transporter | 5,849 ± 442 |
| *Pp*-CIFR-8 #10 | 1,826,219 | 1627 | + | hypothetical protein | 6,185 ± 311 |
| *Pp*-CIFR-8 #11 | 1,794,153 or 3,818,388 | 1599 or 3373 | - | *bamA-I* or *bamA-II*, outer membrane protein assembly factors | 5,732 ± 585 |

| | | | | | |
|---|---|---|---|---|---|
| *Pp*-CIFR-8 #12 | 5,937,809 | 5207 | + | *rbbA*, ribosome-associated ATPase | 11,187 ± 497 |
| *Pp*-CIFR-8 #13 | 5,526,697 | 4860 | + | mannose-6-phosphate isomerase | 11,111 ± 744 |
| *Pp*-CIFR-8 #15 | 5,001,555 | 4408 | - | pseudogene | 7,643 ± 510 |
| *Pp*-CIFR-8 #16 | 4,875,297 | 4284 | - | hypotetical transporter | 6,808 ± 1,744 |

After isolation of antibiotic resistant and fluorescent colonies, transposon insertion sites were mapped on the chromosome via arbitrary PCR. The last column reports the normalized fluorescence of the different colonies. The fluorescence intensity values represent average values from 2 independent measurements ± standard deviation.

**Table S6.** Transposon insertion sites of pCIFR in *E. coli* BW25113.

| Colony | Genome coordinate (bp) | BW25113 number | Strand | Gene name, putative function |
|---|---|---|---|---|
| *Ec*-CIFR-2 #1 | 2,775,129 | 2647 | + | *ypjA*, adhesin-like autotransporter YpjA/EhaD |
| *Ec*-CIFR-2 #5 | 57,403 | 0054 | - | *lptD*, lipopolysaccharide assembly protein LptD |
| *Ec*-CIFR-2 #8 | 249,907 | 0234 | - | *yafP*, putative N-acetyltransferase YafP |
| *Ec*-CIFR-2 #10 | 153,461 | 0141-0142 | + | Intergenic region |
| *Ec*-CIFR-2 #11 | 249,881 | 0234-0235 | - | intergenic region |
| *Ec*-CIFR-2 #18 | 4,371,896 | 4154 | - | *frdA*, fumarate reductase flavoprotein subunit |
| *Ec*-CIFR-2 #20 | 1,046,905 | 0990 | + | *cspG*, cold shock protein CspG |
| *Ec*-CIFR-2 #21 | 249,881 | 0234 | - | *yafP*, putative N-acetyltransferase YafP |
| *Ec*-CIFR-2 #22 | 340,799 | 0326 | - | *yahL*, uncharacterized protein YahL |
| *Ec*-CIFR-2 #23 | 2,182,596 | 2109 | - | *yehB*, putative fimbrial usher protein YehB |
| *Ec*-CIFR-2 #24 | 1,218,262 | 1172 | - | *ymgG*, PF13436 family protein YmgG |
| *Ec*-CIFR-2 #28 | 249,847 | 0234-0235 | - | intergenic region |
| *Ec*-CIFR-4 #32 | 2,875,937 | 2759 | - | *casB*, type I-E CRISPR system Cascade subunit CasB |
| *Ec*-CIFR-4 #33 | 3,459,543 | 3335 | + | *yheC*, type II secretion system prepilin peptidase |
| *Ec*-CIFR-4 #38 | 212,674 | 0193 | + | *yaeF*, peptidase C92 family protein YaeF |
| *Ec*-CIFR-4 #40 | 3,067,667 | 2928 | - | *yggC*, P-loop NTPase domain-containing protein YggC |
| *Ec*-CIFR-4 #42 | 4,401,439 | 4183 | - | *yjfK*, putative transporter YifK |
| *Ec*-CIFR-4 #48 | 4,519,217 | 4304 | + | *sgcC*, putative PTS enzyme IIC component SgcC |

: After isolation of Km resistant and fluorescent colonies, transposon insertion sites were mapped on the chromosome via arbitrary PCR.

**Table S7.** Transposon insertion sites of pCIFR in *C. necator* H16.

| Colony | Chromosome | Genome coordinate (bp) | H16 number | Strand | Gene name, putative function |
|---|---|---|---|---|---|
| *Cn*-CIFR-2 #1 | 1 | 1,345,102 | A1240 | + | branched-chain amino acid ABC transporter permease |
| *Cn*-CIFR-2 #4 | 1 | 1,163,289 | A1067 | - | NTP pyrophosphohydrolase |
| *Cn*-CIFR-2 #6 | 2 | 1,807,473 | B1603 | - | putative spermidine/putrescine transport system ATP-binding protein |
| *Cn*-CIFR-2 #8 | 1 | 3,161,266 | A2925 | - | *argH*, argininosuccinate lyase |
| *Cn*-CIFR-2 #9 | 1 | 3,161,267 | A2925 | - | *argH*, argininosuccinate lyase |
| *Cn*-CIFR-2 #10 | 1 | 41,945 | A0027 | + | hypothetical protein |
| *Cn*-CIFR-2 #11 | 1 | 100,238 | A0087 | - | *corA-I* |
| *Cn*-CIFR-2 #12 | 1 | 4,015,087 | A3717 | + | tripartite tricarboxylate transporter substrate-binding protein |
| *Cn*-CIFR-2 #13 | 1 | 2,541,998 | A2342 | - | *hrpA*, ATP-dependent RNA helicase HrpA |
| *Cn*-CIFR-2 #15 | 1 | 865,663 | A0792 | + | *pheA*, prephenate dehydratase |
| *Cn*-CIFR-2 #16 | 2 | 112,531 | B0097 | - | *eutC*, ethanolamine ammonia-lyase subunit EutC |
| *Cn*-CIFR-2 #17 | 1 | 3,483,787 | A3227 | - | hypothetical protein |

| | | | | | |
|---|---|---|---|---|---|
| *Cn*-CIFR-4 #18 | 2 | 112,809 | B0097 | + | *eutC*, ethanolamine ammonia-lyase subunit EutC |
| *Cn*-CIFR-4 #19 | 1 | 1,272,992 | A1171 | - | *recJ*, single-stranded-DNA-specific exonuclease RecJ |
| *Cn*-CIFR-4 #20 | 2 | 2,658,125 | B2348-B2349 | - | intergenic region |
| *Cn*-CIFR-4 #23 | 1 | 452,574 | A0432 | - | nucleotidyltransferase family protein |
| *Cn*-CIFR-4 #27 | 2 | 358,711 | B0320 | + | DeoR/GlpR family DNA-binding transcription regulator |
| *Cn*-CIFR-4 #28 | 1 | 225,563 | A0213 | - | VOC family protein |
| *Cn*-CIFR-2 #29 | pGH1 | 114,350 | PHG116 | + | *kup-III*, potassium transporter Kup |
| *Cn*-CIFR-2 #30 | 1 | 411,784 | A0393 | - | DJ-1/PfpI family protein |
| *Cn*-CIFR-2 #31 | 1 | 1,728,039 | A1594 | - | autotransporter outer membrane beta-barrel domain-containing protein |
| *Cn*-CIFR-2 #37 | 1 | 3,922,985 | A3646 | - | ParA family protein |
| *Cn*-CIFR-2 #38 | 2 | 1,695,072 | B1508 | + | LysR substrate-binding domain-containing protein |
| *Cn*-CIFR-4 #39 | pGH1 | 198,371 | 33100-33105 | - | intergenic region |
| *Cn*-CIFR-4 #40 | 2 | 1,938,230 | B1704 | + | LysR family transcriptional regulator |
| *Cn*-CIFR-4 #41 | pGH1 | 154,573 | PHG153-154 | + | intergenic region |
| *Cn*-CIFR-4 #43 | pGH1 | 189,272 | PHG175 | - | hypothetical protein |

| | | | | | |
|---|---|---|---|---|---|
| *Cn*-CIFR-4 #46 | 1 | 3,023,547 | A2796 | + | cupin domain-containing protein |
| *Cn*-CIFR-4 #47 | 1 | 3,448,532 | A3191 | - | tripartite tricarboxylate transporter substrate-binding protein |

After isolation of Km resistant and fluorescent colonies, transposon insertion sites were mapped on the chromosome via arbitrary PCR.

**Table S8.** Transposon insertion sites of pCIFR harboring chromoprotein expressing cassettes in *P. putida* KT2440.

| Colony | Genome coordinate (bp) | PP number | Strand | Gene name, putative function |
|---|---|---|---|---|
| *Pp*-CIFR-2::$^{fw}$Yellow | 682,125 | 0584 | + | methyl-accepting chemotaxis transducer |
| *Pp*-CIFR-2::$^{spis}$Pink | ? | PP_t05/t35/t51 | ? | tRNA-Ile |
| *Pp*-CIFR-2::$^{ae}$Blue | ? | PP_23SA/B/C/D/E/F/G | ? | 23S ribosomal RNA |
| *Pp*-CIFR-2::$^{gfas}$Purple | ? | PP_23SA/B/C/D/E/F/G | ? | 23S ribosomal RNA |
| *Pp*-CIFR-2::$^{fw}$Yellow+**CIFR-4::$^{fw}$Yellow** | 5,166,412 | 4546 | - | *hrpA*, ATP-dependent helicase |
| *Pp*-CIFR-2::$^{fw}$Yellow+**CIFR-4::$^{spis}$Pink** | 5,292,388 | 4666 | - | *mmsB*, 3-hydroxyisobutyrate dehydrogenase |
| *Pp*-CIFR-2::$^{fw}$Yellow+**CIFR-4::$^{gfas}$Purple** | 572,058 | 0486 | - | GntR family transcriptional regulator |
| *Pp*-CIFR-2::$^{fw}$Yellow+**CIFR-4::$^{ae}$Blue** | 4,929,956 | 4338 | + | *cheA*, chemotaxis histidine kinase CheA |
| *Pp*-CIFR-2::$^{spis}$Pink+**CIFR-4::$^{fw}$Yellow** | 1,129,642 | 0989 | - | *gcvH-I*, glycine cleavage system protein H |
| *Pp*-CIFR-2::$^{spis}$Pink+**CIFR-4::$^{spis}$Pink** | 516,619 | 0429 | + | short-chain dehydrogenase/reductase family oxidoreductase |
| *Pp*-CIFR-2::$^{spis}$Pink+**CIFR-4::$^{gfas}$Purple** | 5,347,987 | 4704 | - | hypothetical protein |
| *Pp*-CIFR-2::$^{ae}$Blue+**CIFR-4::$^{fw}$Yellow** | 119,185 | 0114 | - | *metN*, methionine ABC transporter ATP-binding protein |
| *Pp*-CIFR-2::$^{ae}$Blue+**CIFR-4::$^{spis}$Pink** | ? | PP_23SA/B/C/D/E/F/G | ? | 23S ribosomal RNA |
| *Pp*-CIFR-2::$^{ae}$Blue+**CIFR-4::$^{ae}$Blue** | 5,152,599 | 5696-4537 | + | Intergenic region |

| | | | | |
|---|---|---|---|---|
| *Pp*-CIFR-2:: $^{ae}$Blue+**CIFR-4:: $^{gfas}$Purple** | 4,827,856 | 4243 | - | *pvdL*, non-ribosomal peptide synthetase |
| *Pp*-CIFR-2:: $^{gfas}$Purple+**CIFR-4:: $^{spis}$Pink** | 580,171 | 0492 | - | *fdhE*, formate dehydrogenase formation protein |
| *Pp*-CIFR-2:: $^{gfas}$Purple+**CIFR-4:: $^{ae}$Blue** | 2,487,610 | between 2181-2182 | - | Intergenic region |
| *Pp*-CIFR-2:: $^{gfas}$Purple+**CIFR-4:: $^{gfas}$Purple** | 4,823,713 | 4243 | + | *pvdL*, non-ribosomal peptide synthetase |

After selecting the most intensely colored colony for every population of transconjugant transformed with one or two pCIFRs containing a chromoprotein (CP) expression module, an arbitrary PCR was performed on them and sent for Sanger sequencing. All the strains producing two CPs are derived from one of the first four strains, that produces only one type of CP. For these strains, the second transformation is highlighted in bold, and the positional data are referred to that transformation. A different internal primer was used to selectively amplify the insertion site of the second integration.

**Table S9.** Transposon insertion sites of pCIFR-4 harboring *crtEBI* from *P. anantis* for biosynthesis of lycopene in *P. putida* KT2440.

| Colony | Genome coordinate (bp) | PP number | Strand | Gene name, putative function | Lycopene content ($A_{471}/OD_{600}$) |
|---|---|---|---|---|---|
| L2 | 3,229,249 | 2829 | - | GntR family transcriptional regulator | 0.221 ± 0.018 |
| L3 | 4,309,576 | 3781 | - | oxygen-independent coproporphyrinogen III oxidase family protein | 0.230 ± 0.015 |
| L4 | 2,204,998 | 1949 | + | GMC family oxidoreductase | 0.263 ± 0.032 |
| L6 | 5,061,219 | 4458-4459 | - | Intergenic region | 0.275 ± 0.042 |
| L9 | 3,853,366 | 3400-3401 | + | intergenic region | 0.263 ± 0.016 |
| L18 | 2,120,893 | 1885 | - | hypothetical protein | 0.270 ± 0.053 |
| L22 | 2,541,921 | 2234 | + | AraC family transcriptional regulator | 0.201 ± 0.042 |
| L26 | 1,142,533 | 1002-1003 | - | Intergenic region | 0.336 ± 0.058 |
| L29 | 6,041,297 | 5294 | + | rph, ribonuclease PH | 0.271 ± 0.024 |

After transforming *P. putida* KT2440 with pCIFR-4::CM1, 9 of the reddest colonies were selected, an arbitrary PCR was performed on them and sent for Sanger sequencing.

**Table S10:** Transposon insertion sites of pCIFR-4 harboring *crtYZ* from *P. anantis* for conversion of lycopene to zeaxanthin in *P. putida* KT2440.

| Colony | Genome coordinate (bp) | PP number | Strand | Gene name, putative function | Zeaxanthin content (mg/$g_{cdw}$) |
|---|---|---|---|---|---|
| Z1 | 3,562,588 | 3149 | + | oxidoreductase | 0.940 ± 0.073 |
| Z2 | 100,559 | 0095 | + | hypothetical protein | 3.310 ± 1.063 |

After transforming L26.1 (Pp-CIFR::*CM1* #26) with pCIFR-4::*CM2*, 2 yellow colonies were isolated, an arbitrary PCR was performed on them and sent for Sanger sequencing. An internal primer, binding to the streptomycin resistance cassette, was used to selectively amplify the insertion site of the second integration.
